# T cell dysregulation and remodeling in pediatric obesity and weight loss

**DOI:** 10.64898/2026.05.19.722395

**Authors:** Ceire A. Hay, Samir U. Sayed, Diego A. Espinoza, Montana Knight, Eric D. Abrams, Jose S. Campos Duran, Maté Z. Nagy, Molly A. Nelson, Sydney A. Sheetz, Pooja Gunnala, Elena N.M. Gonzalez, Jarad Beers, Colleen Tewksbury, Joy L. Collins, Noel N. Williams, Robert B. Lindell, Melanie A. Ruffner, Edward M. Behrens, Kristoffel R. Dumon, Elizabeth Parks Prout, Jorge Henao-Mejia, Sarah E. Henrickson

**Author notes:** Corresponding author: Sarah E. Henrickson.

## Abstract

Obesity is a chronic inflammatory disease associated with immune dysregulation. However, alterations in adaptive immune function remain unclear, particularly in the setting of childhood obesity and weight loss. We defined peripheral T cell dysregulation in a cross-sectional cohort of pediatric participants across weight categories and in a longitudinal cohort of adolescents with severe obesity undergoing bariatric surgery. We found increased expression of activation markers (including PD-1 and CD69) in non-naive CD8^+^ T cells whereas non-naive CD4^+^ T cells were skewed towards Tfh, Th17, and mixed Th2/Th17 populations. Consistent with a hyperactive state, T cells had enhanced capacity for inflammatory cytokine production (including IFN-γ and TNF-α), along with enrichment of gene sets associated with cytokine signaling, cell proliferation, and cell death. Notably, these phenotypic, functional, and transcriptional alterations were not fully resolved after bariatric surgery, despite clinically meaningful weight loss. Together, these findings demonstrate that pediatric obesity leads to dysregulation of adaptive immune function with incomplete normalization after weight loss.

**SUMMARY:** The impact of pediatric obesity on immune cell function is not well understood. This study demonstrates that both CD4^+^ and CD8^+^ T cells are dysregulated in children living with obesity and further identifies that this dysregulated state persists following clinically significant weight loss.

## INTRODUCTION

Obesity is a state of chronic low-grade inflammation that impacts every organ in the body (Lister *et al*., 2023; Schleh *et al*., 2023) and is associated with detrimental health effects including increased risk of co-morbidities (e.g., certain cancers(van Kruijsdijk, van der Wall and Visseren, 2009; Taubes, 2012), type 2 diabetes(Abbasi *et al*., 2017), hypertension (Myette and Flynn, 2024), autoimmunity (Held, Sestan and Jelusic, 2023; Cordeiro *et al*., 2024; Hagman *et al*., 2025), and asthma (Lang *et al*., 2018; Hay and Henrickson, 2021)) as well as increased morbidity and mortality with certain upper respiratory viral infections (i.e., influenza(Honce and Schultz-Cherry, 2019) and COVID-19(Hamer *et al*., 2020)). It is well-established that most adolescents with obesity, defined as a body mass index (BMI) ≥ 95^th^ BMI percentile for sex and age for individuals 2-20 years old, remain obese in adulthood (Simmonds *et al*., 2016; Ward *et al*., 2017), thereby increasing the risk of comorbid diseases and impaired adaptive immune responses across their lifespan (Neidich *et al*., 2017). Notably, the prevalence of severe pediatric obesity, defined as ≥ 120% of the sex and age specific 95^th^ BMI percentile (BMIP95) (Wei *et al*., 2020; Hales *et al*., 2022; Ogden, Freedman and Hales, 2023), has tripled over the past 25 years (Skinner *et al*., 2018) with 6% of U.S children affected today (Noiman *et al*., 2024). This worrisome trend is expected to continue amongst children aged 2-19 years-old (Skinner *et al*., 2018; Ogden *et al*., 2020; Noiman *et al*., 2024; Münte *et al*., 2025). Children living with severe obesity are at an increased risk of metabolic and cardiovascular co-morbidities compared to their obese counterparts (Schwimmer, Burwinkle and Varni, 2003; Skinner *et al*., 2015; Rankin *et al*., 2016; Münte *et al*., 2025)). Moreover, medically significant weight loss in the setting of severe obesity is challenging to accomplish, and bariatric surgery remains one of the only treatment options that can yield durable, substantial weight loss and reduce the burden of obesity-associated co-morbidities(O’Brien *et al*., 2010; Olbers *et al*., 2017; Adams, 2019; Inge *et al*., 2019; Kelly *et al*., 2024).

Obesity has pleiotropic effects on the human immune system (Fang, Henao-Mejia and Henrickson, 2020; Vrieling and Stienstra, 2023; Jiang *et al*., 2025), causing significant alterations to the cellular immune landscape, as well as to the metabolomic and cytokine milieus, in the periphery (Gallistl *et al*., 2001; Dixon *et al*., 2004; Breslin *et al*., 2012), and across tissues (e.g., adipose tissue(Hildreth *et al*., 2021, 2023; Stansbury *et al*., 2023; Hernandez *et al*., 2024), lung (Almond *et al*., 2023), and intestine (Khan *et al*., 2021)). To date, few studies have evaluated the impact of obesity on the frequency and function of peripheral adaptive immune cells in human subjects, with a notable dearth of literature in the setting of pediatric populations (Mattos *et al*., 2016; Carolan *et al*., 2015; Calcaterra *et al*., 2020; Pacifico *et al*., 2006; Bekkering *et al*., 2024; Pugh *et al*., 2022; Carolan *et al*., 2014; Łuczyński *et al*., 2015; Saito *et al*., 2025; Rastogi *et al*., 2012; Tobin *et al*., 2017). The few studies that assess conventional T cell phenotypes in pediatric obesity (Carolan *et al*., 2014; Łuczyński *et al*., 2015; Pugh *et al*., 2022; Bekkering *et al*., 2024; Saito *et al*., 2025) demonstrate conflicting results, with some finding no significant differences in the abundance of peripheral CD3^+^ T cells in children with obesity (Bekkering *et al*., 2024; Saito *et al*., 2025), while others find CD3^+^ T cells are reduced in frequency(Lima *et al*., 2024). Moreover, prior work is overwhelmingly focused on CD4^+^ T cells, with variable evidence of increased frequencies of regulatory CD4^+^ T cells (Tregs) (Saito *et al*., 2025), Th1 cells (Pacifico *et al*., 2006), and Th17 cells (Łuczyński *et al*., 2015; Lima *et al*., 2024), although inconsistencies in helper subset definitions make it challenging to synthesize. To our knowledge, there are no studies that define the impact of pediatric obesity on CD8^+^ T cells. However, a few adult studies demonstrated a positive association between leptin, an adipokine increased in obesity (Kiernan and MacIver, 2020), and frequency of peripheral PD-1^+^CD8^+^ T cells (Wang *et al*., 2019). Beyond phenotype, investigations related to conventional peripheral αβT cell function (Pacifico *et al*., 2006; van der Weerd *et al*., 2012; Lima *et al*., 2024), in the setting of either adult (van der Weerd *et al*., 2012) or pediatric (Pacifico *et al*., 2006; Lima *et al*., 2024) obesity are quite limited. Similarly, there are only a few studies suggesting the phenotype and function of unconventional T cells including natural killer T cells (NKT) (Lynch *et al*., 2009), mucosal-associated invariant T cells (MAIT) (Carolan *et al*., 2015; O’Brien *et al*., 2019) and γδ T cells (Mangan *et al*., 2013) are altered by obesity. Together, these data underscore that much remains to be understood about the impact of obesity on the peripheral immune system – especially in pediatric populations (Fang, Henao-Mejia and Henrickson, 2020; Valentine and Nikolajczyk, 2024).

Although the detrimental effects of obesity on the immune system are well established, there are few interventions that yield durable weight loss, thus the impact of previous obesity on the immune system is not well understood. Preliminary investigations related to T cell phenotype and function in the setting of weight loss in adults (Rizk *et al*., 2021; Gihring *et al*., 2023; Taselaar *et al*., 2025; Dai *et al*., 2017; Zhan *et al*., 2017) have demonstrated increased frequencies of CD4^+^ Tregs (Rizk *et al*., 2021) as well reduced capacity for cytokine production by CD4^+^ T cells (Dai *et al*., 2017; Zhan *et al*., 2017) post-bariatric surgery, when compared to pre-surgical samples. However, investigations with multiple follow-up visits (Gihring *et al*., 2023; Taselaar *et al*., 2025) demonstrate conflicting findings related to the durability of obesity-associated shifts in T cell differentiation. To our knowledge, there are no studies that evaluate the impact of medically significant weight loss on peripheral T cells in adolescents living with obesity.

Here, we investigate the impact of obesity and weight loss in an understudied patient population. Through the integrated analysis of multimodal data, we demonstrate that both CD4^+^ and CD8^+^ T cells are hyperactivated in the setting of pediatric obesity. Furthermore, through the interrogation of peripheral blood samples from adolescents before and after bariatric surgery, we discover T cell activation persists despite clinically significant weight loss. Our data provide a comprehensive characterization of a poorly understood, and, in the case of peripheral CD8^+^ T cells, previously unexplored, immune compartment across two clinically distinct cohorts of pediatric obesity.

## RESULTS

### Peripheral immune cell lineages are stable across pediatric BMIs

To investigate the impact of obesity on the peripheral immune system in children, we studied peripheral blood mononuclear cells (PBMCs) from a cohort of children with obesity (O; > 95^th^ BMI percentile) and age-matched healthy controls (HC; 5^th^-85th BMI percentile) **(Fig. 1 A, Table S1)**. While one group of children in this cohort were living with obesity, all participants were otherwise healthy and had no known comorbidities at the time of recruitment, including no need for daily medications, no recent visits to specialist physicians, and no recent acute illnesses. To appropriately assess obesity in our pediatric populations, we use BMIP95 (the 95^th^ percentile of BMI for age and sex), per the recommendation of the Centers for Disease Control and the American Academy of Pediatrics for children with very high BMI values (Wei *et al*., 2020; Hales *et al*., 2022; Ogden, Freedman and Hales, 2023). BMIP95 expands the dynamic range of the BMIP used in childhood growth trajectory charts, thus improving the ability to differentiate participants living with severe obesity (≥ 120% of BMIP95) (Hampl *et al*., 2023; CDC, 2024) **(Fig. 1 B-C, Table S1)**, which includes 9/15 (60%) of the obese participants in this cohort.

**Figure 1.**
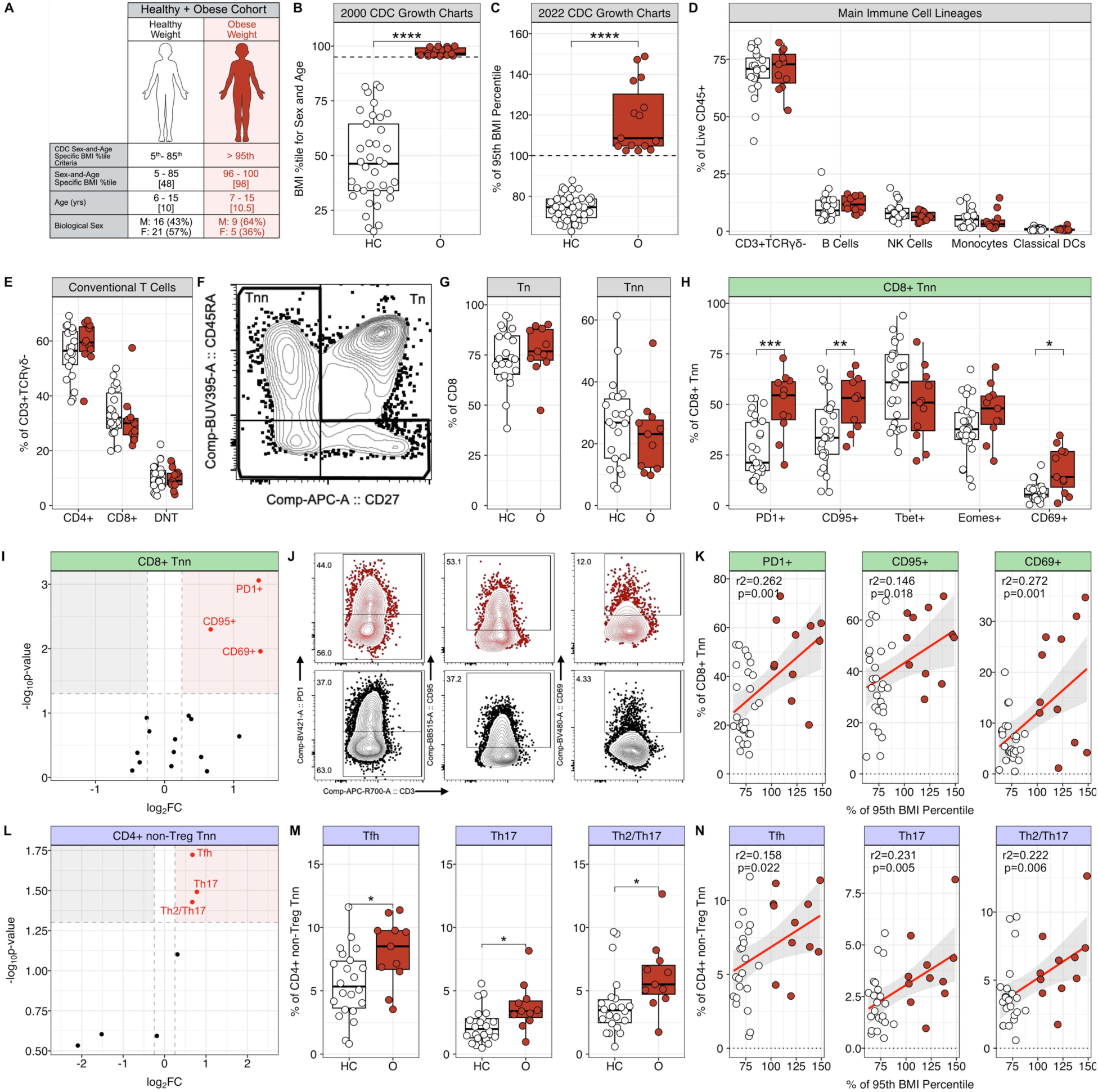
Peripheral T cells from children with obesity exhibit phenotypes consistent with T cell activation. **(A)** Schematic of pediatric cohort. **(B)** Sex-and-Age specific BMI percentile values across Healthy Control (HC, white, n=22) and Obese (O, red, n=11) participants. **(C)** BMI represented as % of 95^th^ BMI percentile (BMIP95) across patient groups. **(D)** Immune lineages in PBMCs expressed as proportion of live CD45^+^ cells. **(E)** T cell subsets expressed as proportion of CD3^+^TCRγο-T cells. **(F)** Representative flow plot of CD8^+^ naive (Tn; CD45RA^+^CD27^+^) and non-naive (Tnn; CD45RA^+/-^CD27^+/-^) in a HC participant. **(G)** Frequency of CD8^+^ Tn and Tnn cells within CD3^+^TCRγδ^-^CD8^+^ T cells. **(H)** Frequency of indicated populations within CD8+ Tnn. **(I)** Volcano plot of log2FC of median frequency of surface markers and transcription factors **(Table S4)** in CD8^+^ Tnn compartment. **(J)** Representative flow plots of PD-1, CD95 or CD69 within CD8^+^ Tnn cells. **(K)** Linear model of frequency of expression of indicated protein within CD8^+^ Tnn compartment vs BMIP95. **(L)** Volcano plot of log2FC of T helper (Th) cell frequencies calculated as median proportion of cells within the Tnn compartment CD4^+^ non-Tregs. **(M)** Frequencies of indicated Th subsets shown as percent of CD4^+^ non-Treg Tnn**. (N)** Linear model of indicated cell Th frequencies within CD4^+^ non-Treg Tnn cells vs BMIP95. For both volcano plots (**I,L**), red and gray background colors highlight proteins with increased frequency of expression in O and HC, respectively. Indicated in red text are proteins significantly increased over HC (p < 0.05) and log2FC ≥ 0.25. For all boxplots and scatterplots, circles represent individual participants. HC and O participants compared by pairwise t tests in boxplots. In all scatterplots (**K,N**), regression line is indicated in red, 95% and confidence interval in gray, and p values are reported for the interaction term between population frequency and BMIP95. Statistical significance reported as ∗ = p < 0.05, ∗∗ = p < 0.01, ∗∗∗ = p < 0.001. For all representative gating (**F,J**), data are represented as 5% contour plots, HC in black and O in red.

To better understand the impact of obesity on the frequency and function of the peripheral immune landscape in pediatric populations, we evaluated immune cell lineages from PBMCs collected from HC (n=22) and O (n=11) donors via a 35-color general immunophenotyping spectral flow cytometry panel (Duran *et al*., 2024) **(Table S2-S3)**. The frequencies of major immune cell subsets were not different between HC and O **(Fig. 1 D, Fig. S1 A-D)**. Within conventional T cells, the proportion of CD4^+^, CD8^+^, and double negative T cells (CD4^-^CD8^-^; DNT) cells were not different between groups **(Fig. 1E)**. Together, these data suggest pediatric obesity does not drive alterations in the balance of either innate or adaptive cells in the periphery.

### Peripheral CD8^+^ cells are skewed towards activated phenotypes in children with obesity

Previous work has suggested an altered immune phenotype in adults with obesity, specifically increased frequency of PD-1^+^ CD8^+^ T cells (McQuade *et al*., 2018; Wang *et al*., 2019). Therefore, we used a 26-color spectral flow cytometry panel focused on T cell activation and effector function (Duran *et al*., 2024) **(Table S4)** to test whether pediatric obesity would demonstrate an increase in expression of inhibitory receptors and activation markers in peripheral CD8^+^ T cells. There were not significant changes in the differentiation state (e.g., central memory, effector memory, TEMRA, naive) of CD8^+^ T cells between HC and O **(Fig. 1 F-G, Fig. S1 E)**. Within the non-naive (Tnn) CD8^+^ compartment, we found increased frequencies of PD-1^+^, CD95^+^, and CD69**^+^** cells **(Fig. 1 H)**, with the median frequencies (PD-1^+^ and CD69^+^ CD8^+^ Tnn cells) demonstrating a more than 4-fold enrichment in the periphery of O compared to HC **(Fig. 1 I)**. The abundances of other single-positive CD8^+^ Tnn populations defined using expression of inhibitory receptors (i.e., CD39, Lag3), activation markers (e.g., CD38, CD49d, HLA-DR, Granzyme-B, and Ki-67), and transcription factors associated with memory (e.g., T-bet) and exhaustion (e.g., TOX, CTLA4, Eomes, Helios, and TCF)(McLane, Abdel-Hakeem and Wherry, 2015; Baessler and Vignali, 2024) were not significantly different between HC and O **(Fig. 1 H-I, Fig. S1 F-H)**. We next sought to determine whether the frequencies of CD8+ Tnn populations demonstrated a linear association with BMIP95. We found that the increased frequencies of PD-1^+^, CD95^+^, and CD69^+^ CD8^+^ Tnn cells **(Fig. 1 J)**, as well as HLA-DR^+^ CD8^+^ Tnn cells, were associated with increased BMIP95 values **(Fig. 1 K, Fig. S1 I)**. Similarly, we used a linear model to interrogate the association between population frequencies and age and found that none of the phenotyping markers **(Table S4)** evaluated in our panel showed a significant association with age **(Fig. S1 J).** Together, these data suggest that the expression of some T cell activation markers increases in a BMIP95-dependent, and not an age-dependent, manner. Moreover, the lack of significant differences across several inhibitory receptors associated with T cell exhaustion, and TOX, a transcription factor critical for promoting a terminally exhausted CD8+ T cell program(Khan *et al*., 2019), suggest that pediatric obesity does not promote an exhausted state in peripheral CD8^+^ T cells. On the contrary, the findings suggest CD8^+^ T cells are skewed towards phenotypes consistent with T cell activation. Given these findings, we next asked whether other adaptive immune cells were similarly altered by pediatric obesity.

### Effector CD4^+^ T cells are skewed towards Tfh and Th17 phenotypes in pediatric obesity

Given the conflicting evidence related to frequencies of CD4^+^ T cell phenotypes in children with obesity(Pacifico *et al*., 2006; Łuczyński *et al*., 2015; Lima *et al*., 2024; Saito *et al*., 2025), we asked whether peripheral CD4^+^ T cells from participants with obesity would demonstrate alterations in differentiation status or subset balance via spectral flow cytometry **(Table S2-S3)**. We gated CD4^+^ T cell subsets **(Fig. S1 K)** and found no significant differences in the frequency of either Tregs (CD4+CD25+CD127-) or non-Tregs (CD4^+^CD25^-^CD127^+/-^) **(Fig. S1 K,L)**, nor were there significant differences in the differentiation state of non-Tregs between HC and O **(Fig. S1 K,M)**. To identify T helper subsets, we evaluated chemokine receptor (e.g. CXCR5, CXCR3, CCR4, CCR6) expression within Tnn compartment of CD4^+^ non-Tregs **(Fig. S1 K).** We found the proportion of follicular helper (Tfh; CD4^+^/CD25^+/-^CD127^+/-^/Tnn/CXCR5^+^), Th17 (CXCR5^-^/CCR4^-/^CCR6^+^CXCR3^-^), and Th2/Th17 (CXCR5^-^/CCR4^+^/CCR6^+^CXCR3^-^) cells were increased in the CD4+ Tnn compartment of O compared to HC **(Fig. 1 L-M, Fig. S1 K)**. We highlight that while the proportions of these subsets were significantly increased in O **(Fig. 1 M)**, their enrichment **(Fig. 1 L)** was less striking (<1.4-fold enrichment) than the alterations we observed in the CD8+ Tnn compartment **(Fig. 1 I)**. We did not observe significant differences in Th1 (CXCR5^-^/CCR4^-^/CC4R6-CXCR3^+^), Th2 (CXCR5^-^/CCR4^+^/CCR6^-^CXCR3^+^), Th1/Th2 (CXCR5^-^/CCR4^+^/CCR6^-^CXCR3^+^, and Th1/Th17 (CXCR5^-^/CCR4^-^/CCR6^+^CXCR3^+^) subsets **(Fig. S1 N)**. Across CD4^+^ T helper subsets, we found increased frequencies of Tfh, Th17, Th2/17, as well as Th2 cells, were associated with increased BMIP95 values **(Fig. 1 N, Fig. S1 O).** In contrast, the frequency of Th1/Th17 cells were positively associated with age **(Fig. S1 P)**. Together, these data suggest the balance of peripheral T helper subsets frequencies are driven primarily by BMIP95-dependent, and not age-dependent, differences in HC and O participants.

### The obese CD4^+^ T cell transcriptome is enriched for pathways associated with effector functions

To investigate the underlying mechanisms by which pediatric obesity alters the peripheral immune landscape, we purified live cells via fluorescence-activated cell sorting (FACS) from PBMCs of HC and O **(Table S5)** for cellular indexing of transcriptomes and epitopes by sequencing (CITE-seq), a transcriptional sequencing assay that allows for the simultaneous assessment of transcripts and surface protein levels via antibody dependent tags (ADT) (Stoeckius *et al*., 2017). We first identified key immune cell lineages (e.g., CD4^+^ T cells, CD8^+^ T cells, monocytes, B cells, NK cells) within this PBMC data before isolating CD4^+^ T cells for transcriptome-based clustering. Here, we identified 24 unique clusters of CD4^+^ T cells **(Fig. S2 A)** which were then assigned to 6 transcriptionally related groups (CD4^+^ T naive, CD4^+^ T helper, CD4^+^ Treg, CD4^+^ Tγδ, CD4^+^ T cycling) **(Fig. 2 A, Table S6)** based on transcript **(Fig. 2 B, Fig. S2 B, Table S7)** and ADT **(Fig. S2 C, Table S8)** expression. There were no differences in cluster abundance between HC and O across unique clusters **(Table S9)** or cluster groups **(Table S10)**. We next sought to determine if transcriptional states were altered by pediatric obesity. To test this, we performed pseudobulk differential gene expression analysis (differentially expressed genes in **Table S11-S12**), followed by gene set enrichment analysis (GSEA)(Subramanian *et al*., 2005) using Hallmark gene sets (Liberzon *et al*., 2015), focusing on the CD4^+^ T helper cluster group given our previous data demonstrating altered proportions of T helper subsets **(Fig. 1 L-M)**. We found pathways related to cytokine signaling and inflammation (e.g., TNFA signaling via NF-κB, IL-2 STAT5 Signaling), cytokine production (e.g. Myogenesis) (Kwee *et al*., 2018), cell proliferation (e.g., G2M Checkpoint, Mitotic Spindle, P53 Pathway), and cell death (e.g., Apoptosis) were enriched in O participants **(Fig. 2 C, Table S13)**. In HC, we found enrichment of pathways associated with cell metabolism (e.g., Oxidative Phosphorylation, Fatty Acid Metabolism, Peroxisome) and cell growth (e.g. Myc Targets v1) **(Fig. 2 C, Table S13)**. We found similar results across the remaining CD4^+^ T clusters groups **(Fig. S2 D, Table S13)**. These results suggest pathways related to cytokine production and inflammation are activated in the peripheral CD4^+^ T cells of children living with obesity.

**Figure 2.**
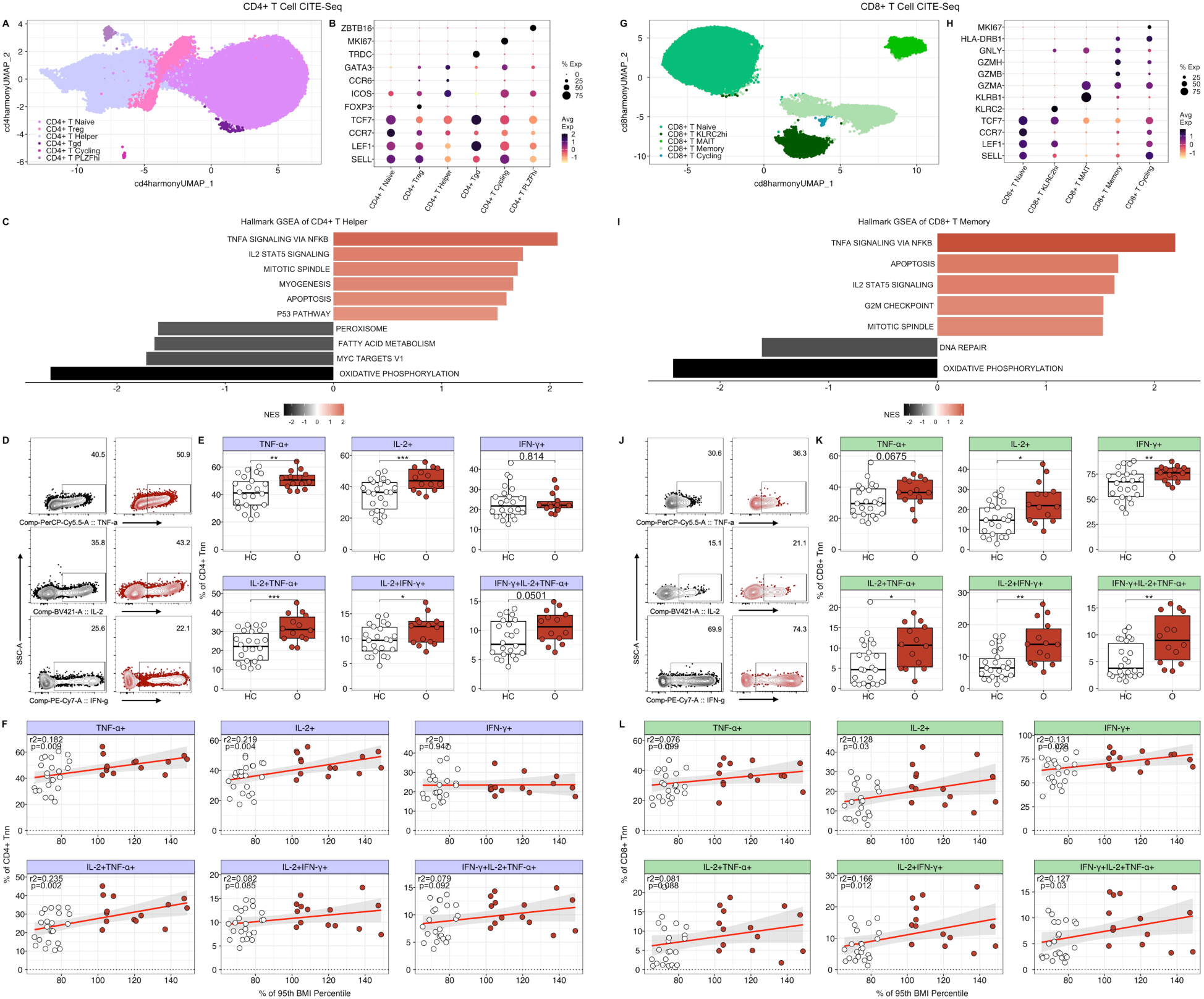
Peripheral T cells from obese participants demonstrate increased capacity for cytokine production. **(A)** UMAP plot of CITE-seq CD4^+^ T cells with six transcriptionally distinct groups. **(B)** Bubble plot of transcripts used to distinguish CD4+ T cell clusters. **(C)** Bar plot of hallmark GSEA results from the CD4^+^ T helper cluster group. **(D)** Representative flow cytometry gating for intracellular cytokine production within CD4^+^ Tnn cells after stimulation with PMA/Ionomycin. **(E)** Quantification of frequency of CD4^+^ Tnn cells that produce indicated cytokine. **(F)** Linear model of intracellular cytokine frequencies within CD4^+^ Tnn cells and BMIP95. **(G)** UMAP plot of CITE-seq CD8^+^ T cells showing represented as 5 transcriptionally distinct groups. **(H)** Bubble plot of transcripts used to distinguish CD8^+^ T cell clusters. **(I)** Bar plot of hallmark GSEA results derived from CD8^+^ T memory cluster group. **(J)** Representative flow cytometry gating for intracellular cytokine production within CD8^+^ Tnn cells. **(K)** Quantification of frequency of CD8^+^ Tnn cells that produce indicated cytokine. **(L)** Linear model of intracellular cytokine frequencies within CD8^+^ Tnn cells and BMIP95. For bubble plots **(A,G)**, color scheme reflects change in average gene expression in O vs. HC (scaled log-normalized counts) ranging from -1 (yellow) to 2 (black) and size of circle reflects proportion of cells expressing indicated transcript. For bar plots **(C,F),** positive NES values indicate pathways upregulated in O (shown in red) and negative NES values represent pathways upregulated in HC (shown in black). Representative gating **(D,J)** is shown as 5% contour plots with HC in black and O in red. In all scatterplots **(F,L),** regression line is indicated in red, 95% and confidence interval in gray, and p values are reported for the interaction term between population frequency and BMIP95. For all boxplots **(E,K)**, HC and O compared by pairwise t-tests in boxplots. Statistical significance reported as ∗ = p < 0.05, ∗∗ = p < 0.01, ∗∗∗ = p < 0.001. For all boxplots **(E,K)** and scatterplots **(F,L)**, circles represent individual participants

### Increased capacity for inflammatory cytokine production by peripheral CD4^+^ T cells in pediatric obesity

Given the significant enrichment of cytokine signaling pathways in CD4^+^ T helper cells, we next used PMA/Ionomycin stimulation to measure the capacity for cytokine production by CD4^+^ Tnn cells **(Table S14)**. We found increased frequencies of TNF-α^+^ and IL-2^+^ cells **(Fig 2. D, Table S15)**, as well as increased frequencies of IL-2^+^TNF-α^+^ and IL-2 IFN-γ^+^ cells **(Fig. 2 E, Fig. S2 E, Table S15),** in O vs HC. We next asked whether the frequencies of cytokine-producing CD4^+^ Tnn cells were linearly associated with BMIP95. We found that increased frequencies of IL-2^+^TNF-α^+^ cells were associated with increased BMIP95 values **(Fig. 2 F, Fig. S 2F)**. In contrast, we did not find an association with age across the cytokine populations we evaluated (**Fig. S2 G).** These results suggest the enhanced capacity for proinflammatory cytokine production in the CD4^+^ Tnn cells of children with obesity is BMIP95-depedent. Together with our transcriptional analysis, this data suggests pediatric obesity drives activation of peripheral CD4^+^ T cells.

### The obese CD8^+^ T cell transcriptome is enriched for pathways associated with effector functions

To better understand the transcriptomic changes driven by obesity in peripheral CD8^+^ T cells, we isolated CD8+ T cells from our CITE-seq data and preformed transcriptome-based clustering to identify 16 unique clusters **(Fig. S2 H)**, which we then assigned to one of 5 transcriptionally related groups (CD8^+^ T naive, CD8^+^ T KLRC2^hi^, CD8^+^ T MAIT, CD8 T memory, CD8^+^ T cycling) **(Fig. 2 G, Table S16)** based on both transcript **(Fig. 2 H Fig. S2 I, Table S17)** and ADT **(Fig. S2 J, Table S18)** expression. There were no differences in cluster abundance between HC and O across unique clusters **(Table S19)** or cluster groups **(Table S20)**. As with our CD4^+^ T cell analysis, we next preformed pseudobulk differential gene expression analysis (differentially expressed genes in **Table S22-22**) followed by GSEA using Hallmark gene sets, focusing here on the CD8+ T cell memory cluster group given our previous data demonstrating that obesity drives an altered phenotype in CD8^+^ Tnn cells **(Fig. 1)**. Using this approach, we found pathways related to cytokine signaling and inflammation (e.g., TNFA signaling via NF-κB, IL-2 STAT5 Signaling), cell proliferation (e.g., G2M Checkpoint, Mitotic Spindle), and cell death (e.g., Apoptosis) were enriched in O whereas DNA Repair and Oxidative Phosphorylation pathways were enriched in HC **(Fig. 2 I, Table S23)**. We found similar results across grouped CD8^+^ T cell clusters **(Fig. S2 K, Table S23).** In line with our findings in CD4^+^ T cells, the enrichment for pathways associated with cytokine production and inflammation suggests the transcriptome of peripheral CD8+ T cells is skewed towards inflammatory gene expression in children living with obesity.

### Increased capacity for inflammatory cytokine production by peripheral CD8^+^ T cells in pediatric obesity

Given the significant enrichment of cytokine signaling pathways in CD8^+^ T memory cells, we next examined the capacity of CD8^+^ Tnn cells from HC and O to produce effector cytokines following *ex vivo* simulation with PMA/Ionomycin **(Table S14)**. We found increased proportions of CD8^+^ Tnn cells producing effector cytokines both individually (TNF-α^+^, IL-2^+^, IFN-γ^+^) **(Fig. 2 J-K, Table S15)** and in combination (IL-2^+^TNF-α^+^, IL-2^+^IFN-γ^+,^ IL-2^+^TNF-α^+^IFN-γ^+^) **(Fig. 2 K, Fig. S2 L, Table S15)** in O vs HC. Furthermore, we found the increased frequencies of TNF-α^+^ **(Fig. 2 L)** and IFN-γ^+^TNF-α^+^ **(Fig. S2 M)** CD8^+^ Tnn cells were associated with increased BMIP95. Within CD8^+^ Tnn cells, we highlight the increased frequencies of several cytokine populations (e.g., IL-2^+^TNF-α^+^, IL-2^+^, TNF-α^+^, IFN-γ^+^TNF-α^+^) were linearly associated with increased age **(Fig. S2 N)**. These results suggest, in contrast to our CD4^+^ T cell data, the increased capacity for proinflammatory cytokine production by CD8^+^ Tnn is impacted by both BMIP95 and age. Taken together with our transcriptomic data **(Fig. 2 C)**, these findings support the hypothesis that peripheral CD8^+^ T cells are skewed towards a hyperactivated state in children living with obesity.

### Bariatric surgery induces weight loss in adolescents with severe obesity

Having demonstrated phenotypic, transcriptional, and functional alterations driven by pediatric obesity in peripheral T cells, we hypothesized weight loss would ameliorate at least some of the alterations we identified in the CD4^+^ T cell **(Fig. 1 L-N, Fig. 2 left)** and CD8^+^ T cell **(Fig. 1 H-K, Fig. 2 right)** compartments of O vs HC. To test this, we collected and studied PBMCs from 11 adolescent participants before and at a median 8.8 ± 2.8 months after undergoing a robotic sleeve laparoscopic gastrectomy (Seeras, Sankararaman and Lopez, 2025), a bariatric surgery procedure in which part of the stomach is removed to facilitate weight loss in individuals living with severe obesity **(Fig. 3 A, Table S24)**. We also collected PBMCs from age-matched controls **(Fig. 3 A, Table S24)**. We highlight that bariatric surgery participants are living with significantly higher BMIs pre-operatively than participants from our first cohort **(Fig. 3 B)**, with 11 of 11 (100%) participants meeting severe obesity criteria **(Fig. 3 C left, Table S24)** and 6 of 11 participants living with at least one additional chronic condition **(Table S25)**. Within this cohort, weight loss outcomes were as expected(O’Brien *et al*., 2010; Olbers *et al*., 2017; Inge *et al*., 2019; Kelly *et al*., 2024), with participants losing an average of 28.6 kg (± 9.7) representing a mean 23 (± 6.4) % reduction from baseline weight **(Fig. 3 C)**. While all patients included in this study lost at least 10% of their baseline weight **(Fig. 3 D)**, we note that 7 of 11 (64%) participants still met criteria for obesity **(Fig. 3 C)** at the post-surgical timepoint (8.8 ± 2.8 months). This outcome reflects that successful surgical outcomes amongst adolescents with severe obesity leave participants with an improved, but not resolved, degree of obesity.

**Figure 3.**
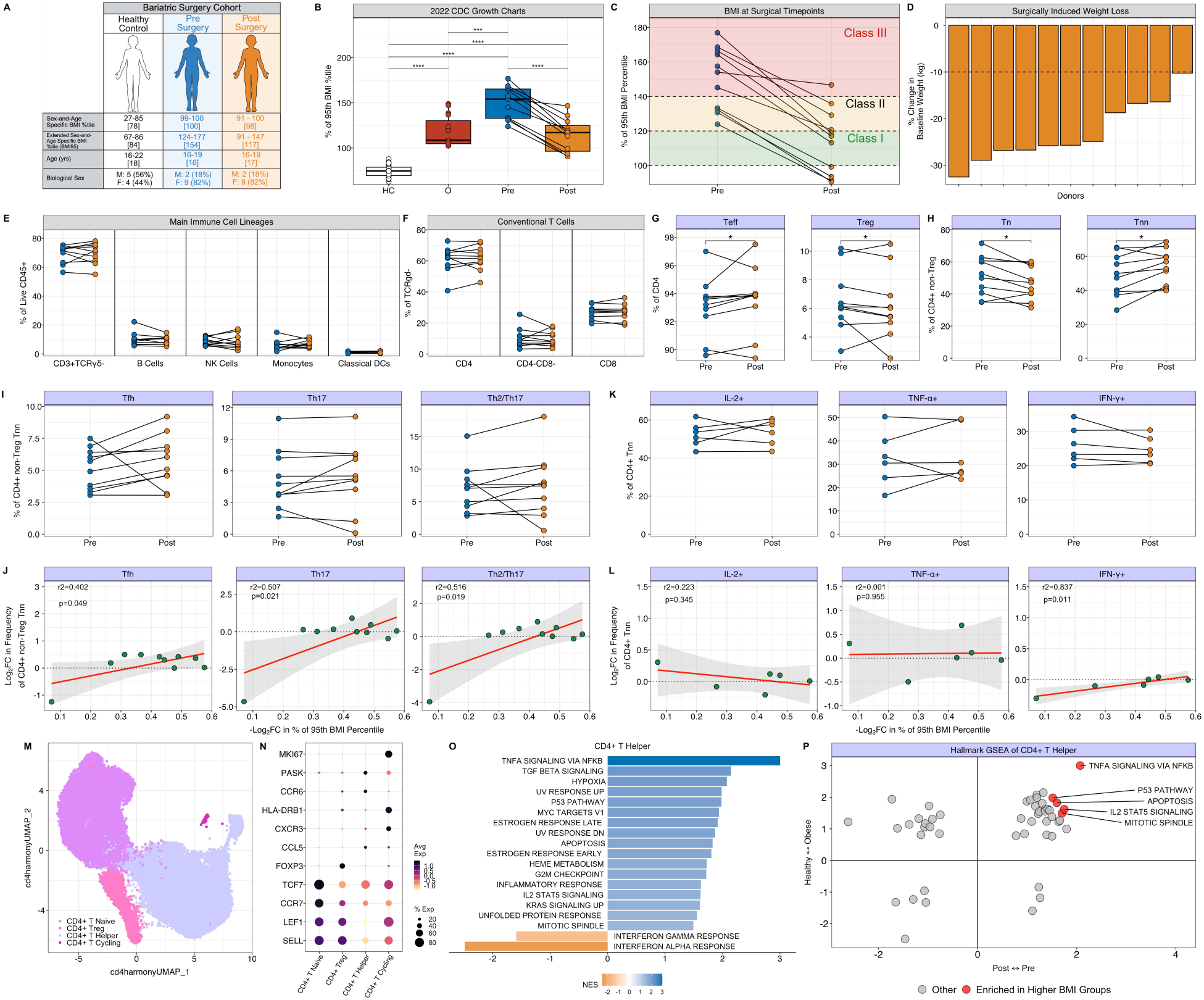
Weight loss alters the peripheral CD4^+^ T cell compartment in adolescents. **(A)** Schematic of bariatric surgery cohort and age-matched controls. **(B)** Quantification of BMI values expressed as BMIP95 adjusted for age and sex across HC (red), O (red), Pre (blue), and Post bariatric surgery (orange). **(C)** BMIP95 values across paired Pre and Post bariatric surgery participants. Green, yellow, and red shading represent criteria for class I, class II, and class III obesity, respectively. **(D)** Bar plot with percent change in weight post-operatively for individual participants with dotted line indicating 10% weight loss. **(E)** Quantification of indicated immune cell populations plotted as percentage of live CD45^+^ cells, **(F)** percentage of CD3^+^TCRγδ^-^ T cells, **(G)** percentage of CD4^+^ T cells and (H) percentage of CD4^+^ Teff cells across paired participants. **(I)** Quantification of indicated subsets show as % of CD4^+^ non-Treg Tnn. **(J)** Scatter plot demonstrating negative log2FC in weight loss compared to log2FC in frequency of indicated population within CD4^+^ non-Treg Tnn compartment. Negative log2FC in weight loss was calculated by dividing post-surgical BMIP95 by pre-surgical BMIP95 and multiplying the log2 value of the result by -1 such that larger absolute value indicates increased weight loss. Log2FC in population frequency was determined by dividing the cell frequency at post-surgical timepoint (Pre) by pre-surgical timepoint (Post) and calculating the log2 value of the result. **(K)** Quantification of proportion of CD4^+^ Tnn cells that produce indicated cytokines after PMA/ionomycin stimulation. **(L)** Scatterplot of the log2FC in frequency of indicated cytokines within CD4^+^ Tnn cells shown compared to -log2FC in BMIP95 calculated as described above. **(M)** UMAP plot of CITE-seq CD4^+^ T cells representing 4 transcriptionally similar groups. **(N)** Bubble plot of transcripts used to distinguish CD4^+^ T cell clusters. The color scheme reflects z-score values within indicated cluster ranging from -1 (yellow) to 2 (black) and size reflects proportion of cells expressing indicated transcript. **(O)** Bar plot of hallmark GSEA results from analysis of the CD4^+^ T helper cluster. Blue bars represent pathways enriched in pre-surgery participants and orange represents pathways enriched in post-bariatric surgery participants. **(P)** Quadrant plots indicating Hallmark GSEA NES from grouped CD4^+^ T helper clusters across HC vs O and Pre vs Post analyses. Circles represent unique Hallmark gene sets, with red circles indicating pathways significantly enriched in both O (vs HC) and Pre (vs Post), and gray circles representing the pathways that are not significantly enriched in both groups. For boxplot and line plots, circles represent individual participants with paired participants indicated by a line. Statistical analysis via pairwise t-test except between paired participants which were compared via paired t-test. For scatterplots, regression line is indicated in red, 95% and confidence interval in gray, and p values are reported for the interaction term between log2FC in frequency and -log2FC in BMIP95. Statistical significance reported as ∗ = p < 0.05, ∗∗ = p < 0.01, ∗∗∗ = p < 0.001.

### Peripheral immune cell lineages are not altered by bariatric surgery-induced weight loss

To assess the impact of weight loss in the setting of severe obesity on immune cell frequency and function, we used spectral flow cytometry **(Table S2-3)** to evaluate immune cell lineages from the PBMCs of adolescent participants (n=11) at paired timepoints before and after bariatric surgery. We found proportions of broad categories of immune cells (e.g., CD3^+^ T cells, B cells, NK cells, monocytes, dendritic cells) were not significantly different in frequency between pre-bariatric surgery (Pre) and post-bariatric surgery (Post) participants **(Fig. 3 E, Table S24)**. While we found γδT cells were decreased in frequency in Post vs Pre **(Fig. S3 A)**, frequencies of conventional T cell subsets **(Fig. 3 F)** and other subsets of innate immune cells **(Fig. S3 A-C)** were not significantly different between Pre and Post samples. Overall, these data suggest the abundance of most peripheral immune cell lineages are not altered by weight loss.

### Weight loss does not alter proportions of peripheral CD4^+^ T helper populations

Given that previous studies that have shown increased proportions of Tregs in the periphery of adults following bariatric surgery-induced weight loss (Rizk *et al*., 2021), we next used spectral flow cytometry to evaluate CD4^+^ T cells from PBMCs of adolescents before and after bariatric surgery. Within CD3^+^TCRγδ^−^CD4^+^ T cells, we identified decreased proportions of CD4^+^ non-Treg **(Fig. 3 G left)** and increased proportions of CD4+ Tregs **(Fig. 3 G right)** in Post vs Pre. Moreover, cells within the CD4^+^ non-Treg compartment were skewed towards a Tnn phenotype in Post vs Pre **(Fig. 3 I)**, although the frequencies of individual memory subsets (i.e., Tcm/Tem/TEMRA) were not significantly different between timepoints **(Fig. S3 D)**. Next, we hypothesized that the frequencies of Tfh, Th17, and Th2/Th17 within the Tnn compartment of non-Tregs, which we had shown were increased in O vs HC **(Fig. 1 L-M)**, would be decreased following weight loss. However, we instead found the frequencies of Tfh, Th17, and Th2/Th17 subsets **(Fig. 3 I),** as well as the remaining T helper subsets (Th1, Th2, Th1/Th2, Th1/Th17) **((Fig. S3 E)**, were not significantly different between Pre and Post. To understand if weight loss led to alterations of the CD4^+^ T cell compartment that were consistent with healthy weight adolescents, we also compared post-surgical samples to aged-matched healthy controls (HC) **(Fig. S3 F)**. We did not detect differences in Th subset frequences between HC and Post **(Fig. S3 F)**, suggesting Th subset frequencies are within normal limits following weight loss. Next, because Tfh, Th17, and Th2/Th17 subsets were trending towards increased frequencies in Post vs Pre **(Fig. 3 I)**, we sought to evaluate if the change in population frequencies between surgical timepoints was linearly associated with change in weight between surgical timepoints. To accomplish this, we calculated the log2 fold-change (log2FC) of T helper frequencies at post timepoints (Post) divided by pre timepoints (Pre) and calculated weight change as the negative log_2_FC (-log2FC) in BMIP95 of Post divided by Pre. We then applied a linear model with these parameters and found the increased log2FC of Tfh, Th17, and Th2/Th17 frequencies were associated with increased weight loss **(Fig. 3 J, Fig. S3 G).** These results suggest that the frequencies of these helper subsets increased, rather than decreased, in a weight-loss dependent manner following bariatric surgery. Finally, we sought to determine if the change in T helper proportions was associated with elapsed time since surgery. To test this, we calculated the log2FC in T helper frequencies (described above) as well as the difference in participant age in months between Post and Pre timepoints. After applying a linear model with these parameters, we did not find a significant relationship between change in frequency and time since surgery across the T helper subsets we evaluated **(Fig. S3 H),** though important to note that we did not evaluate any timepoints later than 12.2 months and our median timepoint was 8.8 months. Taken together, these data suggest that the obesity-associated alterations we found in the CD4^+^ T cell compartment are not impacted by weight loss in the setting of severe obesity.

### Weight loss does not alter the cytokine production capacity of peripheral CD4^+^ T cells

Given the increased capacity for cytokine production by CD4^+^ Tnn cells we observed in O vs HC **(Fig. 2 D-F)**, we hypothesized cytokine production would be decreased following medically significant weight loss in adolescents with severe obesity. To test this, we stimulated PBMCs derived from adolescents at matched pre- and post-bariatric surgery timepoints with PMA/ionomycin and measured intracellular cytokine production by CD4^+^ Tnn cells **(Table S14)**. The populations frequencies we found to be increased in the CD4^+^ Tnn compartment of O vs HC (e.g., TNF-α^+^, IL-2^+^, IL-2^+^TNF-α^+^, IL-2^+^IFN-γ^+^)**(Fig. 2 H)** were not significantly different between Pre and Post samples **(Fig. 3 K, Fig. S3 I)**, nor were the remaining combinations of proinflammatory cytokines we considered **(Fig. S3 I).** Next, we compared the cytokine production capacity of CD4^+^ Tnn cells in Post samples to age-matched HC. We found the proportion of CD4^+^ Tnn IL-2^+^ cells were increased in Post samples compared to HC **(Fig. S3 J),** suggesting that a hyperactivated peripheral CD4^+^ T cell state persists in the setting of weight loss. To evaluate if the change in proportion of cytokine-producing CD4^+^ Tnn cells between Pre and Post samples was linearly associated with weight loss, we calculated the log2FC in population frequencies (e.g., TNF-α^+^, IL-2^+^, etc.) within the CD4^+^ Tnn compartment and the -log2FC in BMIP95 between paired Post and Pre samples. We found that only the increased log2FC in IFN -γ^+^ CD4^+^ Tnn frequency was associated with increasing weight loss **(Fig. S3 L, Fig. S3 K)**. These data suggest the overall increase in IFN -γ^+^ CD4^+^ Tnn frequency relative to baseline scales with weight loss. Moreover, the log2FC in frequency across all cytokine-producing CD4^+^ Tnn cells we considered in our analysis were not associated with months since surgery **(Fig. S3 L)**. Together, these data suggest the enhanced capacity for proinflammatory cytokine production by CD4^+^ Tnn cells observed in pediatric obesity are not affected by weight loss in adolescents with severe obesity.

### Weight loss induces transcriptional changes in peripheral CD4^+^ T cells

To better understand the effect of weight loss on the transcriptional landscape of peripheral CD4^+^ T cells, we purified live cells from PBMCs via FACS from matched Pre (n=4) and Post (n=4) bariatric surgery participants **(Table S26)** and performed CITE-seq. Using the same methodology described above and in Methods, we first identified key immune cell lineages (e.g., CD3^+^ T cells, monocytes, B cells, NK cells) before isolating CD4^+^ T cells. Here, we performed transcriptome-based clustering to identify 23 unique CD4^+^ T cell clusters **(Fig. S3 M),** which were then categorized into 4 transcriptionally related groups (CD4^+^ T naive, CD4^+^ T helper, CD4^+^ T cycling, CD4^+^ Treg) **(Fig. 3 M, Table S27)** based on RNA **(Fig. 3 N, Fig. S3 N, Table S28)** and ADT **(Fig. S3 O, Table 29)** expression. In contrast to our HC vs O data, we found significant differences in cluster abundances between groups: 4 of 23 clusters (Naive 1, Naive 2, Naive SOX4hi, Memory ICOS^hi^ 1) **(Fig S3 P, Table 30)** were less abundant in Post vs Pre, and 5 of 23 clusters (Naive 4, Memory CXCR5^hi^, Memory ICOS^hi^ 2, Memory CD95^hi^, Memory CCL5^hi^) were more abundant in Post vs Pre **(Fig S3 P, Table 30)**. Amongst cluster groups, CD4^+^ T naive were decreased in abundance whereas CD4^+^ T helper were increased in abundance in Post vs Pre **(Fig. S3 Q, Table 31),** in agreement with our findings of decreased CD4^+^ Tn frequencies in Post vs Pre samples via spectral flow cytometry **(Fig. 3H).** We next performed pseudobulk differential gene expression analysis and found, in stark contrast to our HC vs O analysis **(Table S11-S12)**, 466 differentially expressed genes (DEGs) across all Pre vs Post CD4^+^ T cell comparisons **(Table S32-S33)**. We highlight that in 3 of 4 unique clusters less abundant in Post vs Pre (Naive 1, Naive 2, Naive SOX4^hi^), ZFP36, a gene encoding an RNA-binding protein shown to negatively regulate IL-2, IFN-γ, and TNF-α^+^ production in T cells via mRNA decay (Ogilvie *et al*., 2005, 2009; Moore *et al*., 2018; Salerno *et al*., 2018; Cook *et al*., 2022; Petkau *et al*., 2022), demonstrated the greatest reduction in Post vs Pre **(Fig. S3 R, Table S32).** Additionally, genes associated with NF-κB pathway (e.g. NFKBIA) were decreased in Naive SOX4^hi^ and Memory ICOS1 in Post vs Pre. Overall, these data suggest genes potentially critical for the regulation of inflammatory cytokine production are decreased in peripheral CD4^+^ T cells of adolescents with severe obesity after weight loss.

To better understand the transcriptional impact of weight loss on peripheral CD4^+^ T cells, we preformed GSEA using Hallmark gene sets **(Fig. S3 S, Table S34)**. Across cluster groups, we found that pathways related to cytokine signaling and inflammation (e.g., TNFA Signaling via NF-κB Signaling, TGFB Signaling), cytokine production (e.g., Inflammatory Response), cell proliferation (e.g., G2M Checkpoint, Mitotic Spindle, P53 Pathway), and cell death (e.g., Apoptosis) were enriched in paired Pre vs Post samples **(Fig. S3 S, Table S34)**. In contrast, we found enrichment of the Interferon Gamma Response and Interferon Alpha Response pathways in paired Post vs Pre samples **(Fig. S3 S, Table S34)**. These data suggest that while transcriptional features related to cytokine production are decreased following weight loss, transcriptional features related to interferon signaling pathways are activated following weight loss.

To directly compare the transcriptional alterations within the peripheral CD4^+^ T cells across our distinct cohorts **(Fig. 1 A, Fig. 3 A)**, we next compared the normalized enrichment scores (NES) derived from CD4^+^ T helper cluster groups from HC vs O **(Fig. 2 C)** and Pre vs Post **(Fig. 3 O)** analyses. We found shared enrichment for gene sets associated with cytokine signaling (e.g., TNFA Signaling via NF-κB Signaling, IL-2 STAT5 Signaling), cell proliferation (e.g., Mitotic Spindle), and cell death (e.g., Apoptosis), in Pre (vs Post) and in O (vs HC) **(Fig. 3 P)**. Of these pathways, we note that TNFA Signaling via NF-κB Signaling demonstrated the greatest enrichment across both high BMI groups (e.g., Pre and O) **(Fig. 3 P)**. The shared enrichment of these pathways suggests that similar transcriptional programs are altered within the CD4^+^ T cells of these clinically distinct settings of obesity. Moreover, these findings imply that the obesity-driven transcriptional programs of CD4^+^ T helper cells are modulated by weight loss.

### Expression of activation markers and inhibitory receptors are reduced in the CD8^+^ Tnn compartment following bariatric surgery

Given the increased frequency of activation markers we observed within the CD8^+^ Tnn compartment of O vs HC **(Fig. 1 I-K),** we hypothesized that bariatric-surgery induced weight loss would reverse phenotypes consistent with T cell activation within the CD8^+^ Tnn compartment. To test this, we again used a 26-color spectral flow cytometry panel focused on T cell activation and effector function (Duran *et al*., 2024) **(Table S4)** to evaluate PBMC-derived CD8^+^ T cells from matched Pre and Post bariatric surgery participants (n=11). In contrast to our findings in CD4^+^ T cells, weight loss did not alter the differentiation status of CD8^+^ T cells **(Fig. S4 A).** Of the activation markers we found to be significantly increased in O vs HC (PD-1, CD69, CD95) (**Fig 1H),** only the frequency of CD8^+^ Tnn CD69^+^ cells were significantly decreased in Post vs Pre samples **(Fig. 4 A-B)**. However, many of the activation markers and inhibitory receptors that were unchanged in HC vs O (i.e., CD57, Granzyme B, HLA-DR, Ki-67, Tbet, TOX) **(Fig. S2 F-H)**, were decreased in Post vs Pre **(Fig. 4 B-C),** with an approximately 1.4 fold decrease in median frequency of Ki-67^+^ and CD69^+^ CD8^+^ Tnn cells in Post vs Pre **(Fig. 4 C)**. These data suggest that some while markers associated with T cell activation are decreased following medically significant weight loss, most are distinct from the activation markers associated with pediatric obesity **(Fig. 1 H)**. We next compared Post samples to aged-matched HC to better understand if weight loss led to alterations in CD8^+^ Tnn cell compartment consistent with healthy weight adolescents **(Fig. 4D, Fig. S4 D-E)**. We found the frequency of Lag3^+^ cells were significantly increased in Post vs HC **(Fig. S4 D)** whereas the frequency of Tbet^+^ CD8^+^ Tnn cells were decreased in Post vs HC **(Fig. S4 E).** These data suggest that transcription factors and inhibitory receptors typically associated with T cell exhaustion may not have uniform responses to medically significant weight loss. We next sought to determine if the change in proportion of CD8^+^ Tnn populations between Pre and Post samples (**Fig. 4 A-B**) was linearly associated with weight loss. As described above, we applied the log2FC in population frequency and the -log2FC in BMIP95 to a linear model and found that the decreased log2FC of CD8^+^ Tnn CD95^+^ frequency was associated with increasing weight loss (**Fig. 4 E, Fig. S4 F),** implying CD95^+^ CD8^+^ Tnn frequencies decrease in a weight-loss dependent manner. We did not find an association between time since surgery and change in frequency across any of the CD8^+^ Tnn populations we considered **(Fig. S4 G).** Together, these data suggest CD8^+^ Tnn cells are skewed towards a less activated state in post-surgical samples compared to paired pre-surgical samples.

**Figure 4.**
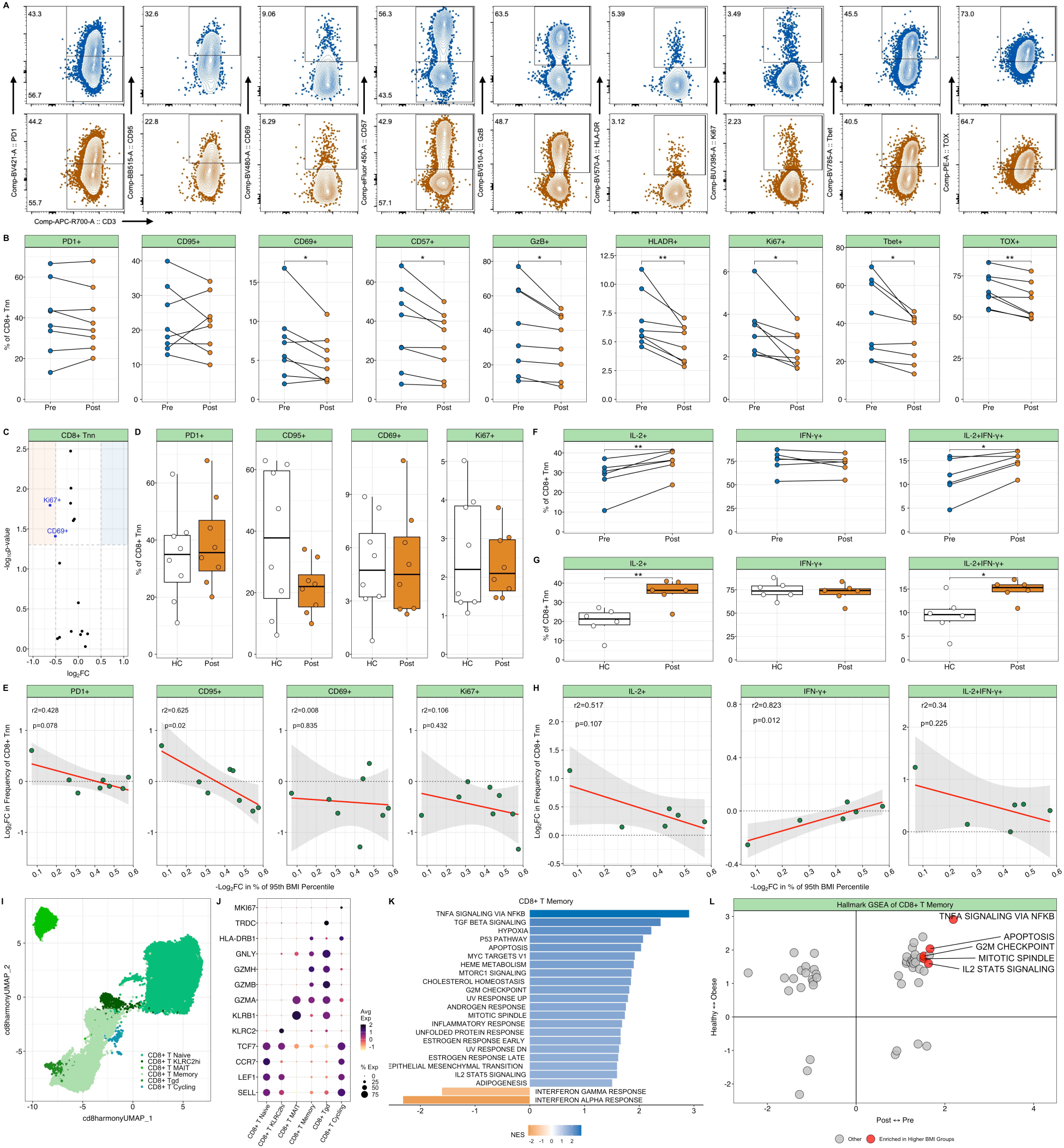
Peripheral CD8^+^ T cells decrease expression of activation markers following weight loss. **(A)** Representative gating of indicated proteins shown as 5% contour plots. **(B)** Quantification of indicated population frequencies within CD8^+^ Tnn. **(C)** Volcano plot of log2FC proteins evaluated via spectral flow cytometry. Blue (upper left) and orange (upper right) colors in the volcano plot highlight proteins elevated in Pre and Post, respectively. Circles represent individual proteins, and blue text indicates proteins significantly downregulated in Post vs Pre. **(D)** Quantification of indicated proteins within CD8^+^ Tnn between HC (white) and Post (orange). **(E)** Scatter plot demonstrating negative log2FC weight loss vs log2FC of frequency of indicated population within CD8^+^ Tnn. **(F,G)** Quantification of indicated cytokine within CD8^+^ Tnn across paired Pre (blue) and Post (orange) **(F)** and between HC (white) and Post **(G)**. **(H)** Scatter plot demonstrating negative log2FC in weight loss compared to log2FC in frequency of indicated cytokine within CD8^+^ Tnn compartment. **(I)** UMAP plot of CITE-seq CD8^+^ T cells representing 6 transcriptionally similar groups. **(J)** Bubble plot of transcripts used to distinguish CD8+ T cell clusters. The color scheme reflects z-score values within indicated cluster ranging from -1 (yellow) to 2 (black) and size reflects proportion of cells expressing indicated transcript. **(K)** Bar plot of hallmark GSEA results from analysis of the CD8^+^ T memory cluster group. Blue bars represent pathways enriched in pre-surgery participants and orange represents pathways enriched in post-bariatric surgery participants. **(L)** Quadrant plots indicating Hallmark GSEA NES from grouped CD8^+^ T memory clusters across HC vs O and Pre vs Post analyses. Dots represent a unique Hallmark gene set and red dots indicate pathways significantly enriched in both O (vs HC) and Pre (vs Post). For boxplot and line plots, circles represent individual participants with paired participants indicated by a line. Statistical analysis via pairwise t test except between paired participants which were compared via paired t-test. For scatterplots, circles represent individual participants, regression line is indicated in red, 95% and confidence interval in gray, and p values are reported for the interaction term between log2FC in frequency and -log2FC in BMIP95. Statistical significance reported as ∗ = p < 0.05, ∗∗ = p < 0.01, ∗∗∗ = p < 0.001.

### Inflammatory cytokine production by CD8^+^ Tnn cells is not altered by weight loss

Given the increased capacity for cytokine production by CD8^+^ Tnn cells we observed in O vs HC **(Fig. 2J-L)**, we hypothesized cytokine production would be decreased following bariatric-surgery induced weight loss. To test this, we stimulated PBMCs derived from adolescents at matched pre-and post-bariatric surgery timepoints with PMA/Ionomycin and measured intracellular cytokine production by CD8^+^ Tnn cells **(Table S14)**. We found increased frequencies of IL-2^+^ and IL-2^+^IFN-γ^+^ CD8^+^ Tnn cells **(Fig. 4 F)**, but not other proinflammatory cytokines **(Fig. S4 H)**, in Post vs Pre samples. Moreover, both IL-2^+^ and IL-2^+^IFN-γ^+^ CD8^+^ Tnn cells were increased in Post vs HC **(Fig. 4 G).** Taken together, these data suggest that despite both significant weight loss and reduction in the frequency of markers associated with activation and effector functions **(Fig. 4 A-E),** the capacity for proinflammatory cytokine production by CD8^+^ Tnn is more pronounced following weight loss. Next, we asked whether the change in proportion of cytokine-producing CD8^+^ Tnn cells (log2FC of Post/Pre) was linearly associated with weight loss (log2FC Post/Pre BMIP95). Of the proinflammatory cytokine populations we considered, only the increased log2FC in IFN-γ^+^ CD8^+^ Tnn cells was associated with increased weight loss **(Fig. 4 H, Fig. S4 J).** These data, which are in line with our findings in CD4^+^ Tnn cells **(Fig. 3 L),** suggests the frequency of IFN-γ producing CD8+ Tnn are increased despite weight loss. We also found that the log2FC TNF-α^+^ CD8^+^ Tnn frequencies were negatively associated with time since surgery **(Fig. S4 K)**. These results suggest peripheral CD8^+^ T cells remain poised towards a proinflammatory state even following clinically significant weight loss, with the production of IL-2 becoming more pronounced in the formerly severely obese state.

### Altered transcriptional landscape of peripheral CD8^+^ T cells in bariatric surgery cohort

To better understand the transcriptomic changes driven by weight loss in peripheral CD8^+^ T cells, we isolated CD8^+^ T cells from our bariatric surgery cohort CITE-seq data **(Table S26)** and performed transcriptome-based clustering to identify 19 discrete CD8^+^ T cell clusters **(Fig. S4 L, Table S35)**, which were then categorized across one of 5 transcriptionally related groups (CD8^+^ T naive, CD8^+^ T KLRC2, CD8^+^ T MAIT, CD8^+^ T memory, and CD8^+^ Tγδ) **(Fig 4 I, Table S35)** based on gene **(Fig. 4 J, Fig. S4 M, Table S36)** and ADT **(Fig. S4 N, Table S37)** expression. Amongst unique cell clusters, we found that the abundance of 2 of 19 clusters (Naive 1 and Memory CD103hi) **(Fig S4 O, Table S38)** were increased in Post samples. Abundances cluster groups did not demonstrate significant changes between Pre and Post samples **(Fig. S4 P, Table S39).** We next preformed pseudobulk differential gene expression analysis and found 280 DEGs between Pre and Post across all CD8^+^ T cell comparisons **(Table S41-42)**. We highlight that clusters enriched in Pre samples (Naive 1 and Memory CD103hi) demonstrated increased expression of ZFP36 **(Fig. S4 Q).** These data, which are in line with our findings in CD4^+^ T cells, similarly suggests genes critical for the regulation of inflammatory cytokine production are altered in the peripheral CD8^+^ T cells of adolescents living with severe obesity.

To better understand the transcriptional impact of weight loss on peripheral CD8^+^ T cells, we preformed GSEA using evaluated Hallmark gene sets **(Fig. S4 R, Table S42)**. Across cluster groups, we that pathways related to cytokine signaling (e.g. TNFA Signaling via NF-κB Signaling, and IL-2 STAT5 signaling), cell proliferation (e.g., G2M Checkpoint, Mitotic Spindle, and P53 Pathway), and cell death (e.g., Apoptosis) were enriched in Pre vs Post **(Fig. S4 S, Table S42)**. We also observed, in agreement with our CD4^+^ T results **(Fig. S3 S)**, enrichment of the Interferon Gamma and Alpha Response pathways in Post vs Pre **(Fig. S4 R, Table S42).** These findings, which are in line with our CD4^+^ T cell data, suggest that peripheral CD8^+^ T cells have transcriptional features consistent with a decreased propensity for cytokine production while also maintaining transcriptional programs associated with inflammatory signal response following medically significant weight loss.

To better understand what pathway-level changes are shared across cohorts, we directly compared NES derived from Hallmark GSEA of CD8^+^ T cell memory cluster groups across the two datasets **(Fig. 2 F, Fig. 4 K)**. Consistent with our CD4^+^ T cell data, we found shared enrichment for TNFA Signaling via NF-κB Signaling, IL-2 STAT5 Signaling, Mitotic Spindle, and Apoptosis pathways in Pre (vs Post) and in O (vs HC) **(Fig. 4 L)**. Similar to our findings in CD4^+^ T helper cells, we note that TNFA signaling via NF-κB demonstrated the greatest enrichment across high BMI groups (e.g., O and Pre) **(Fig. 4 J)**. This shared enrichment for gene sets across high BMI groups suggests, in agreement with our CD4^+^ T helper data, similar transcriptional programs are modulated by these medically distinct settings of obesity and, conversely, by weight loss.

## DISCUSSION

Our study expands upon a limited body of literature that defines immune dysfunction in pediatric obesity (Carolan *et al*., 2014, 2015; Mattos *et al*., 2016; Tobin *et al*., 2017; Bekkering *et al*., 2024). Here, we present a multi-modal characterization of the peripheral adaptive immune system across the spectrum of pediatric obesity, including quantification of altered transcriptional networks across two cohorts that allowed us to define the impact of both obesity and weight loss (in the setting of severe obesity) on immune status in children. We identified BMI-associated increases in the expression of CD69, PD-1, and CD95 in the CD8^+^ Tnn compartment in obese, but otherwise healthy, children. We also demonstrated increased capacity for effector cytokine production, including polyfunctionality, in both CD4^+^ Tnn and CD8^+^ Tnn cells. Furthermore, we interrogated the transcriptional landscape of peripheral T cells across both cohorts using CITE-seq and found shared enrichment of pathways related to cytokine signaling, proliferation, and cell death in the higher BMI groups of both cohorts. To our knowledge, this study provides the first longitudinal characterization of peripheral T cells following weight loss in pediatric subjects, demonstrating that obesity-driven T cell dysregulation persists even after clinically significant weight loss.

Our finding that children living with obesity, in the absence of other medical complications (e.g., asthma, allergy, and metabolic disease), demonstrate increased frequencies of CD95^+^, CD69^+^ and PD-1^+^ cells, within the peripheral CD8^+^ T cell compartment suggests T cells are dysregulated by obesity in childhood. The identification of BMI-associated increases in PD-1^+^ cells is consistent with previous data in adult subjects with obesity in the setting of cancer (Wang *et al*., 2019) and mouse models of obesity (in the setting of cancer (Wang *et al*., 2019; Bader *et al*., 2024) or adipose resident cells (Porsche *et al*., 2021)). This has not been shown in either obese (but otherwise healthy) human subjects or in pediatric subjects broadly. Interestingly, previous work evaluating the impact of pediatric obesity on NK cell frequency and function similarly found increased expression of CD69 and PD-1^58^. Taken together with our findings regarding CD8^+^ Tnn cells, these data suggest that cytotoxic immune cells may be hyperactivated in the setting of pediatric obesity. While there is some data demonstrating altered differentiation states of peripheral T cells following weight loss, a deeper understanding of T cell activation has remained elusive. We highlight a study of bariatric surgery in adults by Villarreal-Calderón, *et. al*., who demonstrated CD69 expression (via MFI of unstimulated cells) was reduced in the PBMCs of adults at post-operative compared to pre-operative timepoints (Villarreal-Calderón *et al*., 2021). This data, taken with our finding of reduced CD69^+^ CD8^+^ Tnn cell frequencies in post-operative adolescent participants, suggest markers of early immune cell activation have detectable decreases in the periphery at when evaluated 8.8 ± 2.8 months after bariatric surgery. We speculate that markers associated with chronic T cell activation or exhaustion (i.e. PD-1, CD39, CTLA-4, Lag3) (Baessler and Vignali, 2024) may demonstrate significant decreases within the peripheral CD8^+^ T cell compartment following prolonged maintenance of weight loss.

Our study provides the potentially novel insight that pediatric obesity augments the capacity for cytokine production by both peripheral CD4^+^ Tnn and CD8^+^ Tnn cells, which is not resolved by clinically meaningful weight loss in adolescent populations. While the impact of weight loss on T cell function has not been rigorously studied in humans broadly (Valentine and Nikolajczyk, 2024), pediatrics remains an understudied patient population (Fang, Henao-Mejia and Henrickson, 2020; Valentine and Nikolajczyk, 2024). Interestingly, peripheral CD4^+^ and CD8^+^ T cells from obese adults have demonstrated defects in activation marker expression (e.g., CD69) and cytokine production (e.g., IFN-γ) following *ex vivo* stimulation with H1N1 (Sheridan *et al*., 2012; Paich *et al*., 2013). Antigen-specific effector function in the context of weight loss has not, to our knowledge, been evaluated in humans, and thus represents a future area of study.

Our finding that memory CD8^+^ and CD4^+^ T helper cells upregulated transcription of Hallmark gene sets related to several cytokine signaling pathways (TNF-α signaling via NF-κB, IL-2 STAT5) in obese subjects (compared to healthy) and pre-operative participants (compared to post-operative) complements previous investigations of adult bariatric surgery. Lo, *et al*. similarly demonstrate enrichment for the TNF-α Signaling via NF-κB and IL-2 STAT5 Signaling pathways at the pre-operative timepoint within a cohort of 6 adults evaluated 3 months post-operatively after bariatric surgery (Lo *et al*., 2021). This study evaluated transcriptomic changes via RNA-sequencing of whole blood, thus T cell data could not be isolated in this analysis. Intriguingly, a recent study of pediatric obesity related-asthma, observed the CD4^+^ T cells of children with obesity alone demonstrated an increased ratio of phosphorylated NF-κB to total NF-κB following *ex vivo* TCR stimulation (Thompson *et al*., 2025). These data, which synergize with our transcriptional findings in CD4^+^ T helper cells, suggests pediatric obesity drives activation of this multipotent transcription factor. We highlight that NF-κB is appreciated as an important mediator of inflammatory disease processes, including in the setting of obesity-associated adipose tissue inflammation^103,104^. As such, we propose the mechanisms by which obesity can promote increased signaling NF-κB signaling peripheral T cells represent an important question for the field.

Obesity is a chronically inflamed state that contributes to significant immune dysregulation (Jiang *et al*., 2025). While the potential mechanisms underpinning the ability of obesity to perturb the immune system remain to be further investigated, here we have identified a hyperactive state of peripheral T cells in pediatric obesity, and further, that significant weight loss is not sufficient to fully resolve this dysregulation in either CD4^+^ or CD8^+^ T cells. Given the persistence of altered T cell phenotype and function across conventional T cell subsets, including altered transcriptional networks, we highlight the investigation of epigenetic remodeling in the setting of pediatric obesity as a potential future direction of this work. There is a growing body of evidence demonstrating obesity epigenetically reprograms murine adipose tissue, resulting in transcriptional and cellular remodeling that persists even after weight loss (Caslin *et al*., 2022; Cottam *et al*., 2022; Hata *et al*., 2023; Hinte *et al*., 2024). These data suggest that the immune system may retain an obesogenic imprint that could maintain immune cells (i.e., CD8^+^ T cells (Cottam *et al*., 2022) and macrophages (Caslin *et al*., 2022; Hata *et al*., 2023)) in a proinflammatory state, thereby contributing to morbidity and mortality disease outcomes in mice (Hata *et al*., 2023). The importance of CD8^+^ T cells in mediating obesity-associated inflammation even after weight loss is underscored in recent work from Garcia*, et. al*., who showed that attenuating memory CD8^+^ T cell responses in murine adipose tissue was protective against metabolic dysfunction associated with weight regain (Garcia *et al*., 2026).

As rates of pediatric obesity continue to rise in the U.S., in parallel with new pharmacologic strategies for weight management (Drucker, 2026), it is imperative that we understand the long term immunological consequences of obesity in children. Equally important is understanding the mechanisms by which weight loss modifies obesity-associated immune dysfunction, which aspects of immune dysfunction can be reversed, and which persist despite successful treatment.

## Study Limitations

The study presented here has several important considerations, which are focused on the bariatric surgery cohort.

1. <u>Preoperative Status:</u> Leading up to bariatric surgery, participants are placed on a modified meal replacement diet to demonstrate weight loss prior to their procedure. Moreover, participants are required to fast overnight prior to surgery. Given that our preoperative samples are collected immediately before surgery, this may not reflect an accurate “baseline” measurement of the subjects’ immune system.
2. <u>Single Post-Surgical Follow-up Timepoint:</u> Bariatric surgery participants were only evaluated at a single timepoint following their surgery, thereby limiting our ability to understand the stability of the changes (or lack thereof) observed compared to their baseline status.
3. <u>Variable Follow-up Timepoint:</u> In addition to being restricted to a single post-operative visit, the post-operative visit was variable across participants (8.8 ± 2.8 months). We selected this this extended time window based on previous data that suggests we can expect a stable weight loss representing 25-30% reduction from baseline within this timeframe (Ryder *et al*., 2024).
4. <u>Complex Comorbidities of Obesity in Surgical Weight Loss Cohort</u>: Unlike participants consented via our standard recruitment protocol, participants from the bariatric surgery cohort are living with co-morbidities including diabetes, asthma, and cardiovascular disease **(Table S25)**. Some participants were eliminated from analysis based on recent surgical history (i.e., organ transplant) or use immunosuppressant drugs (i.e., mTOR inhibitors like everolimus), calcineurin inhibitors (i.e., tacrolimus) and JAK inhibitors (i.e., upadacitinib)). Several participants included in analysis have a history of use of metformin and/or type 2 diabetes as wells as a history of prescription weight loss medication use, either prior to surgery or both before and after their procedure.

## Supporting information

hay_supplemental_tables

hay_supplemental_figures

## MATERIALS AND METHODS

### Ethics Statement

Pediatric healthy donor and bariatric surgery PBMC were collected from participants enrolled in human subjects research with appropriate written consent and approval from the CHOP IRB.

### Human Samples

#### Healthy Control and Obese Cohort

Participants did not have chronic illnesses, daily prescribed medications or recent acute illness.

#### Bariatric Surgery Cohort

Participants had blood samples collected immediately prior to bariatric surgery and again at a follow-up visit 8.8 ± 2.8 months post-surgery. As a comparator, blood samples from age-matched heathy weight participants were also collected. The “ext_bmiz” function from growthcleanr (Lin *et al*., 2022) was used to compute pediatric BMI percentiles, including % of 95^th^ BMI Percentiles, using age in months, height in centimeters, weight in kilograms, and BMI as input.

### Peripheral blood mononuclear cells (PBMCs) Processing

Blood samples were processed by the Children’s Hospital of Philadelphia Biorepository Resource Center Specimen Processing Unit at <6 hours after acquisition. Plasma was separated by centrifugation and snap frozen on dry ice. PBMCs were isolated using SepMate (STEMCELL) density gradient centrifugation using Lymphoprep media (StemCell Technologies; 7861). Whole blood was mixed 1:1 with PBS and layered onto Lymphoprep gradient. SepMate tubes were centrifuged, and the buffy coat suspension was spun down for cell isolation. ACK buffer (Thermo Fisher Scientific; A1049201) was applied to cell pellet to lyse red blood cells and cells were then resuspended in complete RPMI (cRPMI; RPMI 1640 supplemented with 10% FBS, 1% L-Glutamine, 1% PenStrep) for counting prior to cryopreservation in 500μl of freezing media (90% FBS, 10% DMSO). List of reagents available in **Table S43.**

### Spectral Flow Cytometry

PBMCs were plated at 0.25-1x10^6^ cells/well in a 96 well round bottom plate per patient per assay. Cells were washed once with PBS (Penn Cell Center; MT21-031-CM) and stained with an amine-reactive viability dye (Live/Dead Blue, Live/Dead Aqua, or Zombie NIR) in the presence of human Fc block at room temperature (RT) for 15min. Cells were washed with PBS, centrifuged (800G, 2min, RT), and resuspended in 50uL of surface stain prepared in FACS Buffer (PBS + 1% FCS + 2mM EDTA) supplemented with 20% BD Brilliant Stain Buffer (BD Biosciences; 566349). After 30min incubation at 4C, cells were washed once with FACS buffer and resuspended in 50uL of Fix/Perm buffer (Thermo Fisher Scientific; 00-5523-00) to permeabilized cells for intracellular protein staining. After 20min incubation, cells were washed with Perm Buffer (Thermo Fisher Scientific; 00-5523-00), centrifuged, and stained with 50uL intracellular antibody cocktail prepared in Perm Buffer for 1hr at 4C. Following this, cells were washed with Perm Buffer and fixed with 1.6% paraformaldehyde (PFA) (Fisher Scientific; AA433689M) overnight at 4C. The following day, cells were washed once with cold FACS buffer and then resuspended in 125uL of FACS buffer for acquisition Cytek Aurora spectral cytometer (Cytek Biosciences, Fremont, CA). List of antibodies used in flow panels in **(Table S2, S4, S16).** Additional reagents and materials in **Table S43.**

### Intracellular Cytokine Detection

PBMCs were thawed and plated at 0.5x10^6^ cells/well in 96 well round bottom plate. Cells were prepared in cRPMI and plated at 100uL per well. For unstimulated cells, 100uL of cRPMI supplemented with BD GolgiStop (BD Biosciences; 554724) and BD GolgiPlug (BD Biosciences; 555029) was added to wells. For stimulated cells, 100ul of cRPMI containing 100ng/mL PMA (Sigma-Aldrich; P1585-1MG), 1ug/mL Ionomycin (Sigma-Aldrich; I0634-1MG), BD GolgiStop, and BD GolgiPlug was added to wells. After incubation for 4hrs at 37C cells were washed with warm cRPMI and stained for surface and intracellular markers as described above. List of antibodies used in flow panels in **(Table S2, S4, S16).** Additional reagents and materials in **Table S43.**

### Cellular Indexing of Transcriptomes and Epitopes by Sequencing (CITE-seq)

PBMCs were thawed into warm cRPMI, washed once with PBS, and 1x10^6^ cells per donor was aliquoted into separate 5mL FACS tubes. Cells were stained with a 100μL master mix containing Fc block and Zombie NIR in cold PBS. Cells were incubated for 15 minutes at 4°C, washed once with cold cRPMI (500xg for 5 minutes at 4°C). The supernatant was removed and pellets were resuspended with 100μL hashtag oligonucleotides (HTO) (TotalSeq Hashtag Antibodies; BioLegend, Cat. # 394661 – 394976) in cRPMI for 30 minutes at 4°C. Cells were washed twice, resuspended in 100μL cold cRPMI, and passed through a 35μm filter-top 5mL FACS tube. The original tube was rinsed with an additional 100μL cold cRPMI and this volume was added to the filter-top FACS tube. Cells were quick spun, brought up to 400uL with cold cRPMI, and mixed well to prevent cell clumping. Live, single cells were sorted via Cytek Aurora CS System (Cytek Biosciences, Fremont, CA) into 1.5mL Eppendorf tubes containing cold Recovery Buffer (50% FBS in cRPMI with 25mM HEPES). Hash-tagged and sorted cells were centrifuged at 500xg for 5 minutes at 4°C, resuspended in 1mL of cold Recovery Buffer, and recounted. For all CITE-Seq experiments, n=8 unique participants were used to create a pooled sample by aliquoting 100,000 cells per donor into a single 1.5mL Eppendorf. The pooled sample was then incubated with 25μL TruStain FcX blocking reagent (BioLegend, Cat. # 422301) in Cell Staining Buffer (BioLegend, Cat. # 420201) for 10 minutes at 4°C before staining with antibody-derived tags (ADTs) using the TotalSeq-C Universal Cocktail (BioLegend, Cat. # 399905) for an additional 30 minutes at 4°C in a final volume of 50μL. The pooled sample was then washed twice with Cell Staining Buffer and finally resuspended in Cell Staining Buffer at a concentration of 1,500 cells/μL for GEM Generation in a 10X Chromium. List of relevant reagents and materials in **Table S42.**

### 10X GEM Generation, Library Preparation and Sequencing

CITE-seq sample pools were loaded onto 8 wells of a 10X Genomics Chip K (1 pooled sample split across 8 wells) according to the standard protocol for a targeted recovery 5-10,000 cells per well (V2 chemistry). Subsequent steps were performed according to the standard 10X Chromium Next GEM Single Cell 5’ Kit v2 (10x Genomics; Cat. # 1000266) and Chromium Single Cell Human TCR Amplification Kit (10x Genomics; Cat. # 1000252). Prepared libraries were sequenced using an Illumina S2 flow cell (Illumina, San Diego, CA) with a target recovery rate of 24,000 reads/cell for gene expression libraries, 6,000 reads/cell for ADT libraries, and 6,000 reads/cell for TCR libraries. Sequencing was performed at the High Throughout Sequencing Core at the Children’s Hospital of Philadelphia. List of relevant reagents and materials in **Table S42.**

### Bioinformatic Analysis

#### Demultiplexing, mapping and quality control of single cell libraries

Sequencing libraries were demultiplexed using “mkfastq” from 10X Genomics Cell Ranger v7.1.0. Gene expression libraries, ADT libraries, and TCR libraries were then aligned to the human genome (GRCh38), the TotalSeq-C Feature Reference CSV, and the TCR reference (GRCh38), respectively, using 10X Genomics Cell Ranger v7.1.0. Cell Ranger output was then read into Seurat and donors were demultiplexed based on HTO expression via the Seurat function “MULTIseqDemux” (Hao *et al*., 2024). Prior to performing further analysis, we eliminated doublets (as identified by HTOs), dead and dying cells (as identified by more than 12.5% of transcripts originating from mitochondrial genes), and empty droplets (droplets containing fewer than 200 distinct transcripts).

#### Variable feature identification, scaling, dimensionality reduction of single cell data

Samples from each ADT-labelling reaction were merged and ADT expression was normalized within cells using the Seurat function “NormalizeData” (centered log ratio method) prior to performing manual gating and isolation of the CD4^+^ T cell population (utilizing CD3 and CD4 ADT expression) and CD8^+^ T cell population (utilizing CD3 and CD8 ADT expression) to obtain in-silico purified CD4^+^ and CD8^+^ T cell populations. Within experiments, in-silico purified cell populations were then merged for downstream analysis. Following merging, gene expression data was log-normalized and scaled using the Seurat function “LogNormalize” (scale.factor 10000) for each sample. Dimensionality reduction via principal component analysis (PCA) was performed on mean-scaled and centered expression values of commonly-identified variable genes across all samples using the Seurat functions “FindVariableFeatures”, “ScaleData,” and “RunPCA” (Hao *et al*., 2024). PCA reductions were integrated (using sample as the batch variable) by utilizing Harmony. Universal Manifold Approximation and Projections (UMAPs) and nearest-neighbors graphs were constructed using custom code from the *pochi* package (Espinoza, 2022) and clusters were identified from nearest-neighbors graphs using the Seurat function “FindClusters”. Further details and parameters of the analyses are included in the GitHub repository.

#### Annotation of T cells

Clusters were manually annotated based on expression of lineage-defining genes and surface proteins as well as data-defined marker genes and surface proteins. Marker genes were identified utilizing a one-vs-all approach via the “quickMarker” function from the SoupX package(Young and Behjati, 2020), utilizing the term frequency-inverse document frequency (tf-idf) as a measure for marker gene expression. Marker surface proteins (ADTs) were identified via a Wilcoxon test using the “wilcoxauc” function from presto package (Korsunsky *et al*., 2019). We considered ADTs to be predicative of cluster identity when significantly different (p adj < 0.01) and significantly enriched (auc > 0.75). Additional transcripts typically associated with naive, memory, and cycling T cells (e.g., *CCR7, LEF1, SELL, TCF7, IL7R, MK167*) were utilized for further annotation (Zheng *et al*., 2021; Giles *et al*., 2022). Full annotation details are included in the GitHub repository.

#### Gene Set Enrichment Analysis (GSEA)

The function “gseaMultilevel” from the fgsea package (Korotkevich *et al*., 2021) was used to evaluate Hallmark gene sets and we considered gene sets with adjusted p value < 0.01 to be significant.

### Statistical analysis

Statistical analysis was performed utilizing R software via the rstatix package (Kassambara, 2023). Paired T-tests were performed when comparing paired donor samples at discrete timepoints (i.e. pre vs post bariatric surgery). Pairwise T tests were performed when comparing across unique groups of participants (i.e. HC vs O, HC vs Pre, etc.). The R packages dplyr (Wickham *et al*., 2026), tidyr (Wickham, Vaughan and Girlich, 2026), ggplot2(Wickham, 2016), ggpubr (Kassambara, 2026), and cowplot (Wilke, 2025) were utilized for statistical analysis and figure generation.

## Analysis software

All computational analysis was completed using R 4.5.0 and Bioconductor 3.20. Flow cytometry data was processed using FlowJo 10.9.

## Diagrams and Schematics

All schematics were created using BioRender.com.

## Online Supplemental Material

This article contains five supplemental files

1. **Figure S1** shows the analysis of immune cell lineages, including phenotyping of CD4^+^ and CD8^+^ T cells, within the PBMCs of HC and O.
2. **Figure S2** contains the analysis of CITE-seq data and cytokine production capacity data from CD4^+^ T cells (left) and CD8^+^ T cells (right) from the PBMCs of HC and O.
3. **Figure S3** contains the phenotypic, functional, and transcriptional analysis of CD4^+^ T cells from the PBMCs of paired Pre and Post bariatric surgery samples.
4. **Figure S4** shows the phenotypic, functional, and transcriptional analysis of peripheral CD8^+^ T cells from the PMCs of Pre and Post bariatric surgery samples.

This article contains 42 tables

1. **Table S1** provides demographic data for HC and O across all spectral flow cytometry studies.
2. **Table S2** lists the antibody clones used in the general immunophenotyping flow panel.
3. **Table S3** details the gating strategy used to define immune cell lineages in general immunophenotyping panel studies.
4. **Table S4** lists the antibody clones used in the T cell exhaustion flow panel.
5. **Table S5** provides demographic data for HC and O in CITE-seq studies.
6. **Table S6** provides cluster annotations for CD4^+^ T cells derived from CITE-seq analysis of HC and O.
7. **Table S7** contains genes (tf-idf > 0.5) per n=24 unique CD4^+^ T cell cluster from CITE-seq analysis of HC and O.
8. **Table S8** identifies ADTs (auc > 0.75) per n=24 unique CD4^+^ T cell cluster from CITE-seq analysis of HC and O participants.
9. **Table S9** contains the differential abundance analysis results across n=24 unique CD4^+^ T cell clusters from CITE-seq analysis of HC and O.
10. **Table S10** contains the differential abundance analysis results across n=6 CD4^+^ T cell cluster groups from CITE-seq analysis of HC and O.
11. **Table S11** contains the DEGs between HC and O across n=24 unique CD4^+^ T cell clusters
12. **Table S12** contains the DEGs between HC and O across n=6 CD4^+^ T cell cluster groups.
13. **Table S13** provides the Hallmark GSEA results across n=6 CD4^+^ T cell cluster groups between HC and O.
14. **Table S14** lists the antibody clones used in the intracellular cytokine detection flow panel.
15. **Table S15** contains the results of pairwise t-tests for T cell cytokine populations between HC and O participants after stimulation with PMA/ionomycin
16. **Table S16** provides cluster annotations for CD8^+^ T cells derived from CITE-seq analysis of HC and O.
17. **Table S17** contains genes (tf-idf > 0.5) per n=16 unique CD8^+^ T cell cluster from CITE-seq analysis of HC and O.
18. **Table S18** contains ADTs (AUC > 0.75) per n=16 unique CD8^+^ T cell cluster from CITE-seq analysis of HC and O.
19. **Table S19** contains the differential abundance analysis results across n=16 unique CD8^+^ T cell clusters from CITE-seq analysis of HC and O.
20. **Table S20** contains the differential abundance analysis results across n=5 CD8^+^ T cell cluster groups from CITE-seq analysis of HC and O.
21. **Table S21** contains the DEGs between HC and O across n=16 unique CD8^+^ T cell clusters.
22. **Table S22** contains the DEGs between HC and O across n=5 CD8^+^ T cell clusters groups.
23. **Table S23** provides the Hallmark GSEA results across grouped CD8^+^ T cell clusters between HC and O.
24. **Table S24** provides demographic data for bariatric surgery participants in all spectral flow cytometry studies.
25. **Table S25** provides clinical data for bariatric surgery participants in all spectral flow cytometry studies
26. **Table S26** provides demographic data for bariatric surgery participants in CITE-seq studies.
27. **Table S27** provides cluster annotations for CD4^+^ T cells derived from CITE-seq analysis of paired samples from bariatric surgery participants.
28. **Table S28** contains genes (tf-idf > 0.5) per n=23 unique CD4^+^ T cell cluster from CITE-seq analysis of bariatric surgery participants
29. **Table S29** contains ADTs (AUC > 0.75) per n=23 unique CD4^+^ T cell clusters from CITE-seq analysis of bariatric surgery participants.
30. **Table S30** contains the differential abundance analysis results across n=23 CD4^+^ T cell clusters from CITE-seq analysis of bariatric surgery participants.
31. **Table S31** contains the differential abundance analysis results across n=4 CD4^+^ T cell cluster groups CITE-seq analysis of bariatric surgery participants.
32. **Table S32** contains the DEGs between paired samples from bariatric surgery participants across n=23 unique CD4^+^ T cell clusters.
33. **Table S33** contains the DEGs between paired samples from bariatric surgery participants across n=4 CD4^+^ T cell cluster groups.
34. **Table S34** provides the Hallmark GSEA results across n=4 CD4+ T cell cluster groups between bariatric surgery participants.
35. **Table S35** provides cluster annotations for CD8^+^ T cells derived from CITE-seq analysis of bariatric surgery participants.
36. **Table S36** contains genes (tf-idf > 0.5) per n=19 unique CD8^+^ T cell cluster from CITE-seq analysis of Pre and Post samples from bariatric surgery participants.
37. **Table S37** contains ADTs (AUC > 0.75) across n=19 unique CD8^+^ T cell clusters in Pre and Post samples from bariatric surgery participants.
38. **Table S38** contains differential abundance analysis results across n=19 unique CD8^+^ T cell clusters from CITE-seq analysis of Pre and Post samples from bariatric surgery participants
39. **Table S39** contains differential abundance analysis results across n=5 grouped CD8^+^ T cell clusters from CITE-seq analysis of Pre and Post samples from bariatric surgery participants.
40. **Table S40** contains the DEGs between paired Pre and Post samples from bariatric surgery participants across n=19 unique CD8^+^ T cell clusters.
41. **Table S41** contains the DEGs between Pre and Post samples from bariatric surgery participants across n=5 grouped CD8^+^ T cell clusters.
42. **Table S42** provides the Hallmark GSEA results across n=5 grouped CD8^+^ T cell clusters between Pre and Post samples from bariatric surgery participants.
43. **Table S43** lists reagents and materials used in study.

## Data availability

Code and de-identified study data will be made available on a GitHub Repository at the time of publication.

## ACKNOWLEDGEMENTS

First, we would like to thank our patients and their families for participating in this study. We would also like to thank the members of the Children’s Hospital of Philadelphia (CHOP) Biorepository Resource Center Specimen Processing Unit (Richard Tustin III, Annemarie Butler, Vanessa Oliva) for processing all PBMCs related to this project. Finally, we thank the CHOP Flow Cytometry Core members (especially Dr. Florin Tuluc and Jennifer Murray) for their assistance with flow cytometry experiments.

## Funding

University of Pennsylvania Center of Excellence in Environmental Toxicology (CEET)

Translational Research Training Program in Environmental Health Sciences T32 ES019851 (CAH)

University of Pennsylvania Provost’s Graduate Academic Engagement Fellowship at the Netter Center (CAH)

Burroughs Wellcome Fund CAMS (SEH)

CZI Single Cell Analysis of Inflammation Program (JHM and SEH)

Pennsylvania Department of Health 4100089394 (EMB)

NIH/NIGMS K23GM159013 (RBL)

NIH/NIAID K08 AI148456 (MAR)

R01 AI184785 (MAR)

## Declaration of Interests

MAR has received research funding from Regeneron.

## Author contributions

Conceptualization: CAH and SEH

Methodology: CAH and SEH

Investigation: CAH, SS, JSC, and MZN

Data Curation: CAH, EA, SS, JSC, PG, and SEH

Formal Analysis: CAH, DAE, MK, SS, and SEH

Writing – Original

Draft: CAH and SEH

Writing – Reviewing & Editing: all authors

Visualization: CAH, DAE, MK, and SEH

Funding Acquisition: SEH

Supervision: SEH

## Notes

### Competing Interest Statement

The authors have declared no competing interest.

