## Supplementary material for "T cell dysregulation and remodeling in pediatric obesity and weight loss": hay_supplemental_tables: s01_hcob_all_flow_summary.docx

**Table S1**. Demographic data for HC and O across all spectral flow cytometry studies.

| Variable | Healthy N = 37*^a^* | Obese N = 15*^a^* |
| --- | --- | --- |
| Age (years) |  |  |
| Median | 10.00 | 10.00 |
| Min, Max | 6.00, 15.00 | 7.00, 15.00 |
| Sex |  |  |
| Female | 21 (57%) | 6 (40%) |
| Male | 16 (43%) | 9 (60%) |
| Race |  |  |
| Black or African American | 19 (51%) | 11 (73%) |
| Other | 1 (2.7%) | 1 (6.7%) |
| White | 17 (46%) | 3 (20%) |
| BMI Percentile for Sex and Age*^b^* |  |  |
| Median | 46 | 97 |
| Min, Max | 15, 82 | 95, 100 |
| BMI Category |  |  |
| Healthy Weight*^c^* | 37 (100%) | 0 (0%) |
| Obese*^d^* | 0 (0%) | 15 (100%) |
| BMIP95*^e^* |  |  |
| Median | 75 | 109 |
| Min, Max | 63, 88 | 102, 149 |
| Meets Severe Obesity Criteria*^f^* |  |  |
| Yes | 0 (0%) | 6 (40%) |
| No | 37 (100%) | 9 (60%) |
| Severe Obesity Class |  |  |
| Class I Obesity (Non-Severe)*^g^* | 0 (0%) | 9 (60%) |
| Class II Obesity (Severe)*^h^* | 0 (0%) | 4 (27%) |
| Class III Obesity (Severe)*^i^* | 0 (0%) | 2 (13%) |
| N/A | 37 (100%) | 0 (0%) |
| *^a^*n (%) | | |
| *^b^*Used to assess BMI in pediatric populations (2-20 years) | | |
| *^c^*5th-85th BMI Percentile for sex and age | | |
| *^d^*95th ≥ BMI Percentile for sex and age | | |
| *^e^*% of 95th BMI Percentile for sex and age, used for children 2-20 years old with very high BMIs | | |
| *^f^*120% ≥ BMIP95 | | |
| *^g^*100% ≥ BMI95 < 120% | | |
| *^h^*120% ≥ BMI95 < 140% | | |
| *^i^*BMI95 ≥ 140% | | |
