## Supplementary material for "T cell dysregulation and remodeling in pediatric obesity and weight loss": hay_supplemental_tables: s02_gen_panel_abs.docx

**Table S2. Antibodies used in spectral flow general immunophenotyping panel.**

| **N** | **Fluorophore** | **Detector** | **Antigen** | **Clone** | **Vendor** | **Catalog** | **Titration** | **Type** |
| --- | --- | --- | --- | --- | --- | --- | --- | --- |
| 1 | Fc Block | NA | NA | NA | BD Biosciences | 564220 | 200 | Surface |
| 2 | BUV395 | UV2 | CD45RA | H100 | BD Biosciences | 568712 | 200 | Surface |
| 3 | Live/Dead Blue | UV6 | N/A | N/A | Thermo Fisher | L34961 | 1000 | Surface |
| 4 | BUV496 | UV7 | CD16 | 3G8 | BD Biosciences | 612944 | 200 | Surface |
| 5 | BUV563 | UV9 | CD14 | M5E2 | BD Biosciences | 741360 | 200 | Surface |
| 6 | BUV615 | UV10 | CD45 | HI30 | BD Biosciences | 751472 | 400 | Surface |
| 7 | BUV661 | UV11 | CD11c | B-ly6 | BD Biosciences | 612967 | 100 | Surface |
| 8 | BUV737 | UV14 | CD56 | NCAM16.2 | BD Biosciences | 612766 | 200 | Surface |
| 9 | BUV805 | UV16 | CD26 | L272 | BD Biosciences | 749179 | 100 | Surface |
| 10 | BV421 | V1 | PD-1 | EH12.2H7 | BioLegend | 329920 | 100 | Surface |
| 11 | SuperBright 436 | V2 | CD123 | 6H6 | Thermo Fisher | 62-1239-42 | 100 | Surface |
| 12 | eFluor450 | V3 | CD161 | HP-3G10 | Thermo Fisher | 48-1619-42 | 100 | Surface |
| 13 | BV480 | V5 | ICOS | DX29 | BD Biosciences | 746248 | 100 | Surface |
| 14 | BV510 | V7 | Ki67 | B56 | BD Biosciences | 563462 | 50 | ICS |
| 15 | BV570 | V8 | CD3 | UCHT1 | BioLegend | 300436 | 600 | Surface + ICS |
| 16 | BV605 | V10 | CCR4 | L291H4 | BioLegend | 359418 | 100 | Surface |
| 17 | BV650 | V11 | CXCR3 | G025H7 | BioLegend | 353730 | 100 | Surface |
| 18 | BV711 | V13 | CCR6 | G034E3 | BioLegend | 353436 | 100 | Surface |
| 19 | BV750 | V14 | CXCR5 | RF8B2 | BD Biosciences | 747111 | 100 | Surface |
| 20 | BV785 | V15 | CD8 | RPA-T8 | BioLegend | 301046 | 250 | Surface |
| 21 | AF488 | B2 | Foxp3 | 259D | BioLegend | 320212 | 100 | ICS |
| 22 | AF532 | B3 | CD19 | HIB19 | Thermo Fisher | 58-0199-42 | 50 | Surface |
| 23 | PerCP | B8 | CD20 | 2H7 | BioLegend | 302324 | 50 | Surface |
| 24 | PerCP-Cy5.5 | B9 | CD11b | ICRF44 | BioLegend | 301328 | 100 | Surface |
| 25 | PerCP-eFluor710 | B10 | TCRgd | B1.1 | Invitrogen | 46-9959-42 | 150 | Surface |
| 26 | PE | YG1 | IgD | IA6-2 | Biolegend | 348204 | 100 | Surface |
| 27 | cFluor YG584 | YG1 | CD4 | SK3 | CyTEK | R7-20041 | 600 | Surface |
| 28 | PE/Dazzle 594 | YG3 | Tbet | 4B10 | BioLegend | 644828 | 200 | ICS |
| 29 | PE-Cy5 | YG5 | CD25 | M-A251 | BD Biosciences | 555433 | 200 | Surface |
| 30 | PECy7 | YG9 | BCL6 | K112-91 | BD Biosciences | 563582 | 50 | ICS |
| 31 | PE-Fire810 | YG10 | CD39 | A1 | BioLegend | 328245 | 200 | Surface |
| 32 | APC | R1 | CD27 | O323 | BioLegend | 302810 | 100 | Surface |
| 33 | AF647 | R2 | CD33 | P67.6 | BioLegend | 366626 | 200 | Surface |
| 34 | APC-R700 | R4 | CD127 | HIL-7R-M21 | BD Biosciences | 565185 | 100 | Surface |
| 35 | APC-eFluor780 | R7 | CD38 | HIT2 | Thermo Fisher | 47-0389-42 | 100 | Surface |
