## Supplementary material for "T cell dysregulation and remodeling in pediatric obesity and weight loss": hay_supplemental_tables: s03_gen_panel_gating.docx

**Table S3. Gating hierarchy used to define main immune cell lineages and subsets using the general immunophenotyping panel**

| **Full Gating Hierarchy** | **Population Title** |
| --- | --- |
| Cells | Cells |
| Cells/Singlets | Singlets |
| Cells/Singlets/Live CD45+ | Live CD45+ |
| Cells/Singlets/Live CD45+/CD3+ | CD3+ |
| Cells/Singlets/Live CD45+/CD3+/TCRgd+ | gd T cells |
| Cells/Singlets/Live CD45+/CD3+/TCRgd- | ab T cells |
| Cells/Singlets/Live CD45+/CD3+/TCRgd-/CD4 | CD4 |
| Cells/Singlets/Live CD45+/CD3+/TCRgd-/CD4/NonTreg | CD4+ T Effector |
| Cells/Singlets/Live CD45+/CD3+/TCRgd-/CD4/NonTreg/Tcm | CD4+ Tcm |
| Cells/Singlets/Live CD45+/CD3+/TCRgd-/CD4/NonTreg/Tem | CD4+ Tem |
| Cells/Singlets/Live CD45+/CD3+/TCRgd-/CD4/NonTreg/Temra | CD4+ Temra |
| Cells/Singlets/Live CD45+/CD3+/TCRgd-/CD4/NonTreg/Tn | CD4+ Tn |
| Cells/Singlets/Live CD45+/CD3+/TCRgd-/CD4/NonTreg/Tnn | CD4+ Tnn |
| Cells/Singlets/Live CD45+/CD3+/TCRgd-/CD4/NonTreg/Tnn/CXCR5+ | CD4+ Tfh |
| Cells/Singlets/Live CD45+/CD3+/TCRgd-/CD4/NonTreg/Tnn/CXCR5- | CD4+ Non-Tfh |
| Cells/Singlets/Live CD45+/CD3+/TCRgd-/CD4/NonTreg/Tnn/CXCR5-/CCR4+/CCR6+CXCR3- | Th2/17 |
| Cells/Singlets/Live CD45+/CD3+/TCRgd-/CD4/NonTreg/Tnn/CXCR5-/CCR4+/CCR6-CXCR3+ | Th1/2 |
| Cells/Singlets/Live CD45+/CD3+/TCRgd-/CD4/NonTreg/Tnn/CXCR5-/CCR4+/CCR6-CXCR3- | Th2 |
| Cells/Singlets/Live CD45+/CD3+/TCRgd-/CD4/NonTreg/Tnn/CXCR5-/CCR4-/CCR6+CXCR3+ | Th1/17 |
| Cells/Singlets/Live CD45+/CD3+/TCRgd-/CD4/NonTreg/Tnn/CXCR5-/CCR4-/CCR6+CXCR3- | Th17 |
| Cells/Singlets/Live CD45+/CD3+/TCRgd-/CD4/NonTreg/Tnn/CXCR5-/CCR4-/CCR6-CXCR3+ | Th1 |
| Cells/Singlets/Live CD45+/CD3+/TCRgd-/CD4/Treg | CD4+ Treg |
| Cells/Singlets/Live CD45+/CD3+/TCRgd-/CD4-CD8- | DNT |
| Cells/Singlets/Live CD45+/CD3+/TCRgd-/CD8 | CD8 |
| Cells/Singlets/Live CD45+/CD3+/TCRgd-/CD8/Tcm | CD8+ Tcm |
| Cells/Singlets/Live CD45+/CD3+/TCRgd-/CD8/Tem | CD8+ Tem |
| Cells/Singlets/Live CD45+/CD3+/TCRgd-/CD8/Temra | CD8+ Temra |
| Cells/Singlets/Live CD45+/CD3+/TCRgd-/CD8/Tn | CD8+ Tn |
| Cells/Singlets/Live CD45+/CD3+/TCRgd-/CD8/Tnn | CD8+ Tnn |
| Cells/Singlets/Live CD45+/CD3- | CD3- |
| Cells/Singlets/Live CD45+/CD3-/HLADR+/CD20+CD33- | B cells |
| Cells/Singlets/Live CD45+/CD3-/HLADR+/CD20+CD33-/CD11c+Tbet+ | Atypical B cells |
| Cells/Singlets/Live CD45+/CD3-/HLADR+/CD20+CD33-/CD27+IgD+ | Non-Switched B Cells |
| Cells/Singlets/Live CD45+/CD3-/HLADR+/CD20+CD33-/CD27+IgD- | Switched B Cells |
| Cells/Singlets/Live CD45+/CD3-/HLADR+/CD20+CD33-/CD27-IgD+ | Naive B Cells |
| Cells/Singlets/Live CD45+/CD3-/HLADR+/CD20+CD33-/CD27-IgD- | DN B Cells |
| Cells/Singlets/Live CD45+/CD3-/HLADR+/CD20-CD33 | CD20-CD33 |
| Cells/Singlets/Live CD45+/CD3-/HLADR+/CD20-CD33/CD11b+CD14+ | Monocytes |
| Cells/Singlets/Live CD45+/CD3-/HLADR+/CD20-CD33/CD11b+CD14+/CD14++CD16+ | Intermediate Monocytes |
| Cells/Singlets/Live CD45+/CD3-/HLADR+/CD20-CD33/CD11b+CD14+/CD14++CD16++ | Non-Classical Monocytes |
| Cells/Singlets/Live CD45+/CD3-/HLADR+/CD20-CD33/CD11b+CD14+/CD14+CD16- | Classical Monocytes |
| Cells/Singlets/Live CD45+/CD3-/HLADR+/CD20-CD33/CD11b+CD14-/CD11c+CD123- | CD11b+ DCs |
| Cells/Singlets/Live CD45+/CD3-/HLADR+/CD20-CD33/CD11b-CD14-/CD11c+CD123- | Classical Dendritic Cells |
| Cells/Singlets/Live CD45+/CD3-/HLADR+/CD20-CD33/CD11b-CD14-/CD11c-CD123+ | Plasmacytoid Dendritic Cells |
| Cells/Singlets/Live CD45+/CD3-/HLADR-/CD56+ | Natural Killer Cells |
| Cells/Singlets/Live CD45+/CD3-/HLADR-/CD56+/CD56+CD16- | CD56bright NK Cells |
| Cells/Singlets/Live CD45+/CD3-/HLADR-/CD56+/CD56+CD16hi | CD56dim NK Cells |
