## Supplementary material for "T cell dysregulation and remodeling in pediatric obesity and weight loss": hay_supplemental_tables: s04_texh_panel_abs.docx

**Table S4. Antibodies used in spectral flow T cell Exhaustion panel.**

| **N** | **Fluorophore** | **Detector** | **Antigen** | **Clone** | **Vendor** | **Catalog** | **Titration** | **Type** |
| --- | --- | --- | --- | --- | --- | --- | --- | --- |
| 1 | Fc Block | NA | NA | NA | BD Biosciences | 564220 | 200 | Surface |
| 2 | BUV395 | UV2 | Ki67 | B56 | BD Biosciences | 564071 | 300 | ICS |
| 3 | LD Blue | UV6 | Viability | NA | Thermo Fisher | L34962A | 1000 | Surface |
| 4 | BUV496 | UV7 | CD4 | RPA-T4 | BD Biosciences | 741134 | 400+400 | Surface + ICS |
| 5 | BUV563 | UV9 | CD127 | HIL-7R-M21 | BD Biosciences | 748489 | 100 | Surface |
| 6 | BUV737 | UV14 | CD27 | O323 | BD Biosciences | 751681 | 400 | Surface |
| 7 | BUV805 | UV16 | *CD49D+A:B | 9F10 | BD Biosciences | 749454 | 300 | Surface |
| 8 | BV421 | V1 | *PD-1 | EH12.2H7 | BioLegend | 329920 | 100 | Surface |
| 9 | eFluor 450 | V3 | *CD57 | TB01 | Thermo Fisher | 48-0577-42 | 200 | Surface |
| 10 | BV480 | V5 | *CD69 | FN50 | BD Biosciences | 747519 | 200 | Surface |
| 11 | BV510 | V7 | *Granzyme B | GB11 | BD Biosciences | 563388 | 300 | ICS |
| 12 | BV570 | V8 | *HLA-DR | L243 | BioLegend | 307638 | 150 | Surface |
| 13 | BV605 | V10 | CD8a | RPA-T8 | BioLegend | 301040 | 200 | Surface |
| 14 | BV650 | V11 | CD45RA | H100 | BD Biosciences | 563963 | 200 | Surface |
| 15 | BV711 | V13 | *LAG-3 | 11C3C65 | BioLegend | 369320 | 100 | Surface |
| 16 | BV750 | V14 | CCR7 | G043H7 | BioLegend | 353254 | 100 | Surface |
| 17 | BV785 | V15 | *T-bet | 4B10 | BioLegend | 644835 | 300 | ICS |
| 18 | BB515 | B1 | *CD95 | DX2 | BioLegend | 564597 | 100 | Surface |
| 19 | PerCP-Cy5.5 | B9 | KLRG-1 | SA231A2 | BioLegend | 367708 | 100 | Surface |
| 20 | PerCP eFluor710 | B10 | *CD38 | HIT2 | Thermo Fisher | 46-0389-42 | 150 | Surface |
| 21 | PE | YG1 | *TOX | REA473 | Miltenyi | 130-120-716 | 200 | ICS |
| 22 | PE-Dazzle 594 | YG3 | *CD39 | A1 | BioLegend | 328224 | 200 | Surface |
| 23 | PE-Cy5 | YG5 | *CTLA-4 | BNI3 | BD Biosciences | 555854 | 200 | Surface |
| 24 | PE-Cy7 | YG9 | *Helios | 22F6 | Thermo Fisher | 25-9883-42 | 200 | ICS |
| 25 | AF647 | R2 | *TCF1/TCF7 | C63D9 | Cell Signaling Technologies | 6709S | 200 | ICS |
| 26 | APC-R700 | R4 | CD3 | UCHT1 | BD Biosciences | 565119 | 200 | Surface |
| 27 | APC-eFluor780 | R7 | *Eomes | WD1928 | Thermo Fisher | 47-4877-42 | 100 | Surface |
