## Supplementary material for "T cell dysregulation and remodeling in pediatric obesity and weight loss": hay_supplemental_tables: s05_hcob_citeseq_summary.docx

**Table S5.** Demographic data for HC and O in CITE-seq studies.

| Variable | Healthy N = 7*^a^* | Obese N = 8*^a^* |
| --- | --- | --- |
| Age (years) |  |  |
| Median | 10.62 | 9.42 |
| Min, Max | 8.54, 14.15 | 7.98, 13.01 |
| Sex |  |  |
| Female | 6 (86%) | 3 (38%) |
| Male | 1 (14%) | 5 (63%) |
| Race |  |  |
| Black or African American | 2 (33%) | 6 (75%) |
| White | 4 (67%) | 2 (25%) |
| Unknown | 1 | 0 |
| BMI Percentile for Sex and Age*^b^* |  |  |
| Median | 46 | 99 |
| Min, Max | 23, 78 | 96, 100 |
| BMI Category |  |  |
| Healthy | 7 (100%) | 0 (0%) |
| Obese*^c^* | 0 (0%) | 8 (100%) |
| BMIP95*^d^* |  |  |
| Median | 76 | 121 |
| Min, Max | 65, 83 | 103, 149 |
| Meets Severe Obesity Criteria*^e^* |  |  |
| Yes | 0 (0%) | 4 (50%) |
| No | 7 (100%) | 4 (50%) |
| Severe Obesity Class |  |  |
| Class I Obesity (Non-Severe)*^f^* | 0 (0%) | 4 (50%) |
| Class II Obesity (Severe)*^g^* | 0 (0%) | 2 (25%) |
| Class III Obesity (Severe)*^h^* | 0 (0%) | 2 (25%) |
| N/A | 7 (100%) | 0 (0%) |
| *^a^*n (%) | | |
| *^b^*Used to assess BMI in pediatric populations (2-20 years) | | |
| *^c^*95th ≥ BMI Percentile for sex and age | | |
| *^d^*% of 95th BMI Percentile for sex and age, used for children 2-20 years old with very high BMIs | | |
| *^e^*120% ≥ BMIP95 | | |
| *^f^*100% ≥ BMI95 < 120% | | |
| *^g^*120% ≥ BMI95 < 140% | | |
| *^h^*BMI95 ≥ 140% | | |
