## Supplementary material for "T cell dysregulation and remodeling in pediatric obesity and weight loss": hay_supplemental_tables: s14_pma_panel_abs.docx

**Table S14. Antibodies used in spectral flow intracellular cytokine detection panel.**

| N | Fluorophore | Detector | Antigen | Clone | Vendor | Catalog | Titration | Type |
| --- | --- | --- | --- | --- | --- | --- | --- | --- |
| 1 | Fc Block | NA | NA | NA | BD Biosciences | 564220 | 200 | Surface |
| 2 | BUV395 | UV2 | CD4 | RPA-T4 | BD Biosciences | 564724 | 200+ 200 | Surface + ICS |
| 3 | BV421 | V1 | IL-2 | 5344.111 | BD Biosciences | 562914 | 100 | ICS |
| 4 | V500 | V7 | CD14 | M5E2 | BD Biosciences | 561391 | 300 | Surface |
| 5 | V500 | V7 | CD16 | 3G8 | BD Biosciences | 561394 | 300 | Surface |
| 6 | V500 | V7 | CD19 | HIB19 | BD Biosciences | 561121 | 300 | Surface |
| 7 | BV605 | V10 | CD8 | RPA-T8 | BioLegend | 301040 | 200 | Surface |
| 8 | BV650 | V11 | CD45RA | H100 | BD Biosciences | 563963 | 200 | Surface |
| 9 | BV785 | V15 | CD27 | O323 | BioLegend | 302832 | 200 | Surface |
| 10 | AF488 | B2 | IL-17A | BL168 | BioLegend | 512308 | 100 | ICS |
| 11 | PerCP-Cy5.5 | B9 | TNFa | MAb11 | BioLegend | 502926 | 100 | ICS |
| 12 | PE | YG1 | IL-13 | JES10-5A2 | BioLegend | 501903 | 100 | ICS |
| 13 | PE/Dazzle 594 | YG3 | CD39 | 4B10 | BioLegend | 644828 | 200 | Surface |
| 14 | PE-Cy5 | YG5 | CD25 | M-A251 | BD Biosciences | 555433 | 200 | Surface |
| 15 | PE-Cy7 | YG9 | IFNg | B27 | BioLegend | 506518 | 400 | Intracellular |
| 16 | APC-R700 | R4 | CD3 | UCHT1 | BD Biosciences | 565119 | 200 + 200 | Surface + ICS |
| 17 | Zombie NIR | R6 | NA | NA | Thermo Fisher | 423106 | 3000 | Surface |
| 18 | APC-Fire750 | R7 | PD-1 | EH12.2H7 | BioLegend | 329954 | 100 | Surface |
| 19 | APC-Fire810 | R8 | HLA-DR | L243 | BioLegend | 307674 | 200 | Surface |
