## Supplementary material for "T cell dysregulation and remodeling in pediatric obesity and weight loss": hay_supplemental_tables: s24_prepost_all_flow_summary.docx

**Table S24. Demographic summary of bariatric surgery cohort participants used in flow cytometry studies**

| Variable | Healthy Control N = 11*^a^* | Pre N = 11*^a^* | Post N = 11*^a^* |
| --- | --- | --- | --- |
| Age (years) |  |  |  |
| Median | 18.00 | 16.00 | 17.00 |
| Min, Max | 16.00, 23.00 | 16.00, 19.00 | 16.00, 19.00 |
| Sex |  |  |  |
| Female | 5 (45%) | 9 (82%) | 9 (82%) |
| Male | 6 (55%) | 2 (18%) | 2 (18%) |
| Race |  |  |  |
| Black or African American | 6 (55%) | 2 (18%) | 2 (18%) |
| White | 3 (27%) | 6 (55%) | 6 (55%) |
| White and Asian | 1 (9.1%) | 0 (0%) | 0 (0%) |
| Other | 1 (9.1%) | 1 (9.1%) | 1 (9.1%) |
| ND | 0 (0%) | 2 (18%) | 2 (18%) |
| BMI*^b^* |  |  |  |
| Median | 24 | 45 | 35 |
| Min, Max | 20, 26 | 36, 52 | 27, 44 |
| BMI Percentile for Sex and Age*^c^* |  |  |  |
| Median | 78 | 100 | 98 |
| Min, Max | 27, 85 | 99, 100 | 91, 100 |
| Unknown | 3 | 0 | 0 |
| BMI Category |  |  |  |
| Healthy Weight*^d^* | 11 (100%) | 0 (0%) | 0 (0%) |
| Overweight*^e^* | 0 (0%) | 0 (0%) | 4 (36%) |
| Obese*^f^* | 0 (0%) | 11 (100%) | 7 (64%) |
| BMIP95*^g^* |  |  |  |
| Median | 84 | 154 | 117 |
| Min, Max | 67, 86 | 124, 177 | 91, 147 |
| Unknown | 3 | 0 | 0 |
| Meets Severe Obesity Criteria*^h^* |  |  |  |
| Yes | 0 (0%) | 11 (100%) | 4 (36%) |
| No | 8 (73%) | 0 (0%) | 7 (64%) |
| ND | 3 (27%) | 0 (0%) | 0 (0%) |
| Obesity Class |  |  |  |
| Class I Obesity (Non-Severe)*^i^* | 0 (0%) | 0 (0%) | 3 (27%) |
| Class II Obesity (Severe)*^j^* | 0 (0%) | 4 (36%) | 3 (27%) |
| Class III Obesity (Severe)*^k^* | 0 (0%) | 7 (64%) | 1 (9.1%) |
| N/A | 11 (100%) | 0 (0%) | 4 (36%) |
| *^a^*n (%) | | | |
| *^b^*Used to define weight categories in adults 21+ | | | |
| *^c^*Used to assess BMI in pediatric populations (2-20 years) | | | |
| *^d^*5th-85th BMI Percentile for sex and age | | | |
| *^e^*85th-95th BMI Percentile for sex and age | | | |
| *^f^*95th ≥ BMI Percentile for sex and age | | | |
| *^g^*% of 95th BMI Percentile for sex and age, used for children 2-20 years old with very high BMIs | | | |
| *^h^*120% ≥ BMIP95 | | | |
| *^i^*100% ≥ BMI95 < 120% | | | |
| *^j^*120% ≥ BMI95 < 140% | | | |
| *^k^*BMI95 ≥ 140% | | | |
