## Supplementary material for "T cell dysregulation and remodeling in pediatric obesity and weight loss": hay_supplemental_tables: s25_prepost_clinical_summary.docx

**Table S25.** Clinical data for paired bariatric surgery participants in all spectral flow cytometry studies

|  | Clinical Data for Bariatric Surgery Cohort | |
| --- | --- | --- |
| Variable | Pre N = 11*^a^* | Post N = 11*^a^* |
| Diabetes History |  |  |
| Type II Diabetes | 1 (9.1%) | 0 (0%) |
| ND | 10 (91%) | 11 (100%) |
| Other Metabolic Disease |  |  |
| Prediabetes | 2 (18%) | 2 (18%) |
| Insulin Resistance | 4 (36%) | 1 (9.1%) |
| ND | 5 (45%) | 8 (73%) |
| Metformin History |  |  |
| Yes | 0 (0%) | 0 (0%) |
| No | 6 (100%) | 6 (100%) |
| Unknown | 5 | 5 |
| Anti-Obesity Medication |  |  |
| Bupropion/Naltrexone | 1 (9.1%) | 0 (0%) |
| Phentermine Only | 0 (0%) | 1 (9.1%) |
| Topiramate Only | 2 (18%) | 1 (9.1%) |
| Phentermine/Topiramate | 2 (18%) | 1 (9.1%) |
| ND | 6 (55%) | 8 (73%) |
| Asthma History |  |  |
| Current | 1 (9.1%) | 1 (9.1%) |
| Yes | 1 (9.1%) | 0 (0%) |
| ND | 9 (82%) | 10 (91%) |
| Food Allergy History |  |  |
| Current | 2 (18%) | 2 (18%) |
| ND | 9 (82%) | 9 (82%) |
| Drug Allergy History |  |  |
| Current | 2 (18%) | 2 (18%) |
| ND | 9 (82%) | 9 (82%) |
| *^a^*n (%) | | |
