## Supplementary material for "T cell dysregulation and remodeling in pediatric obesity and weight loss": hay_supplemental_tables: s26_prepost_citeseq_summary.docx

**Table S25. Demographic summary of bariatric surgery participants included in CITE-Seq studies.**

| Variable | Pre N = 4*^a^* | Post N = 4*^a^* |
| --- | --- | --- |
| Age (years) |  |  |
| Median | 16.50 | 17.50 |
| Min, Max | 16.00, 18.00 | 16.00, 19.00 |
| Sex |  |  |
| Female | 3 (75%) | 3 (75%) |
| Male | 1 (25%) | 1 (25%) |
| Race |  |  |
| Black or African American | 1 (25%) | 1 (25%) |
| White | 2 (50%) | 2 (50%) |
| ND | 1 (25%) | 1 (25%) |
| BMI*^b^* |  |  |
| Median | 42.9 | 31.2 |
| Min, Max | 38.9, 47.7 | 28.4, 34.8 |
| BMI Percentile for Sex and Age*^c^* |  |  |
| Median | 99.61 | 95.93 |
| Min, Max | 99.12, 99.98 | 90.96, 97.69 |
| BMI Category |  |  |
| Overweight*^d^* | 0 (0%) | 2 (50%) |
| Obese*^e^* | 4 (100%) | 2 (50%) |
| BMIP95*^f^* |  |  |
| Median | 145 | 106 |
| Min, Max | 132, 164 | 91, 117 |
| Meets Severe Obesity Criteria*^g^* |  |  |
| Yes | 4 (100%) | 0 (0%) |
| No | 0 (0%) | 4 (100%) |
| Obesity Class |  |  |
| Class I Obesity (Non-Severe)*^h^* | 0 (0%) | 2 (50%) |
| Class II Obesity (Severe)*^i^* | 2 (50%) | 0 (0%) |
| Class III Obesity (Severe)*^j^* | 2 (50%) | 0 (0%) |
| N/A | 0 (0%) | 2 (50%) |
| *^a^*n (%) | | |
| *^b^*Used to define weight categories in adults 21+ | | |
| *^c^*Used to assess BMI in pediatric populations (2-20 years) | | |
| *^d^*85th-95th BMI Percentile for Sex and Age | | |
| *^e^*95th ≥ BMI Percentile for Sex and Age | | |
| *^f^*% of 95th BMI Percentile for Sex and Age, used for children 2-20 years old with very high BMIs | | |
| *^g^*120% ≥ BMIP95 | | |
| *^h^*100% ≥ BMI95 < 120% | | |
| *^i^*120% ≥ BMI95 < 140% | | |
| *^j^*BMI95 ≥ 140% | | |
