## Supplementary material for "T cell dysregulation and remodeling in pediatric obesity and weight loss": hay_supplemental_figures

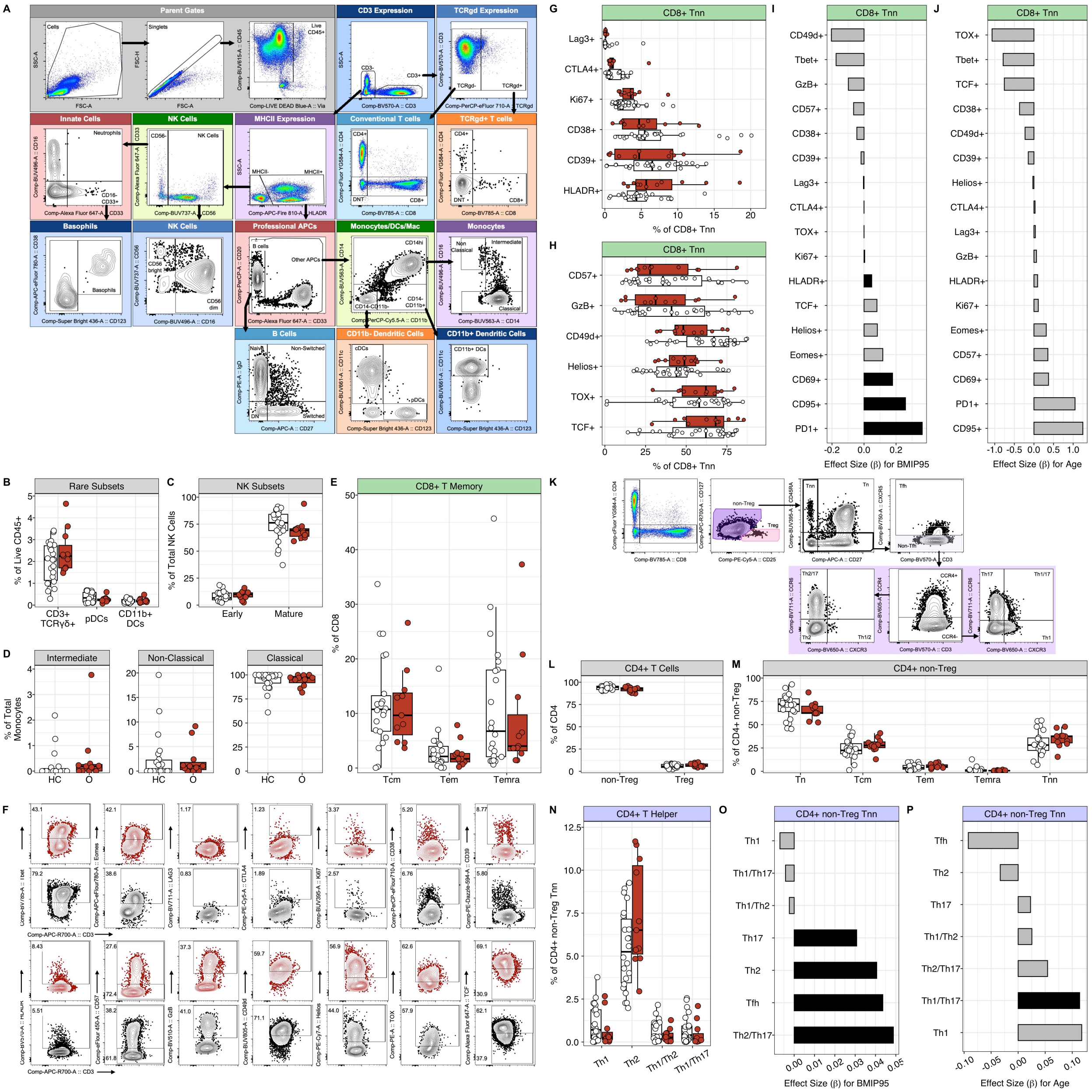

**Supplemental Figure 1. Evaluation of peripheral immune landscape across healthy weight and obese pediatric participants.** Related to Fig. 1. **(A)** Representative flow cytometry plots showing gating strategy to identify major immune cell subsets within a PMBC sample of healthy weight donor. First, cells were identified based on forward (FSC) and side (SSC) properties, then singlets were selected based on FSC height (FSC-H) vs FSC area (FSC-A), followed by selection of live $CD45^+$. T cells were gated based on expression of CD3. $CD3^+\gamma\delta$ T cells and $CD3^+\gamma\delta^-$ T cells (Conventional T cells, Tconv) were then gated based on expression of γδ TCR. Both $CD3^+\gamma\delta$ T cells and Tconv cells were then gated based on expression of CD4 and CD8. From $CD3^-$ gate, cells were further divided into HLA-$DR^+$ and HLA-$DR^-$. Within the HLA-$DR^-$ gate, Natural Killer (NK) cells were defined as $CD56^+CD33^-$ and subsequently divided into $CD56^{bright}$ ($CD56^{bright}CD16^-$) and $CD56^{dim}$ ($CD56^{dim}CD16^+$). Neutrophils were defined within the $MHCII^-CD56^-$ gate as $CD16^+CD33^+$ cells. Basophils were defined within the $CD16^-CD33^+$ gate as $CD123^+CD38^-$ cells. Within the professional APCs gate, B cells were defined as $CD20^+CD33^-$ and were subsequently subdivided into double negative ($CD27^-IgD^-$), naive ($CD27^-IgD^+$), non-switched ($CD27^+IgD^+$), or switched ($CD27^+IgD^-$) B cells. Within the professional APCs gate, other APCs were defined as $CD20^-CD33^+$ and were subsequently divided by expression of CD11b and CD14. The $CD14^{hi}$ gate was used to define monocytes subsets as non-classical ($CD14^-CD16^+$), Intermediate ($CD14^+CD16^{+/-}$), and Classical ($CD14^+CD16^-$) phenotypes. The $CD14^-CD11b^-$ gate was used to define $CD11b^-$ Dendritic Cells (DCs), which were further divided into classical dendritic cells (cDCs; $CD11c^+CD123^-$) and plasmacytoid dendritic cells (pDCs; $CD11c^-CD123^+$) The $CD14^-$ $CD11b^+$ gate was used to define $CD11b^+$ DCs as $CD11c^+CD123^-$ cells. **(B)** Quantification of $TCR\gamma\delta^+$ T cells, pDCs, and $CD11b^+$ DCs cells shown as proportion of live $CD45^+$ cells **(C)** Quantification of Early ($CD16^{bright}$) and Mature ($CD16^{dim}$) NK cells shown as proportion of total NK cells. **(D)** Quantification of non-classical, intermediate, and classical monocytes shown as proportion of total monocytes. **(E)** Quantification of indicated memory subsets within $CD8^+$ T cell compartment. **(F)** Representative gating of indicated markers within the $CD8^+$ Tnn compartment. **(G-H)** Frequency of expression of indicated markers within $CD8^+$ Tnn compartment. **(I)** Bar plot representing the β-coefficients for the BMIP95 term and **(J)** age term in each model. **(K)** Representative flow gating strategy of $CD3^+$ $TCR\gamma\delta^-$ $CD4^+$ peripheral T cells. **(L)** Boxplots quantifying indicated $CD4^+$ T cell subsets. **(M)** Quantification of indicated memory subset frequencies within $CD4^+$ non-Treg compartment. **(N)** Quantification of indicated $CD4^+$ T helper subsets within Tnn compartment of $CD4^+$ non-Tregs. **(O)** Bar plot representing the β-coefficients for the BMIP95 term (Frequency*BMIP95) and **(P)** age term (Frequency*Age) in each model. For all representative gating, data are represented as pseudo color or 5% contour plots where black indicates HC participants. In all bar plots, significant linear model results are represented in black. In boxplots, circles represent individual participants, HC participants indicated in white, O participants indicated in red, and statistical analysis via pairwise t- tests $* = p < 0.05$, $** = p < 0.01$, $*** = p < 0.001$.

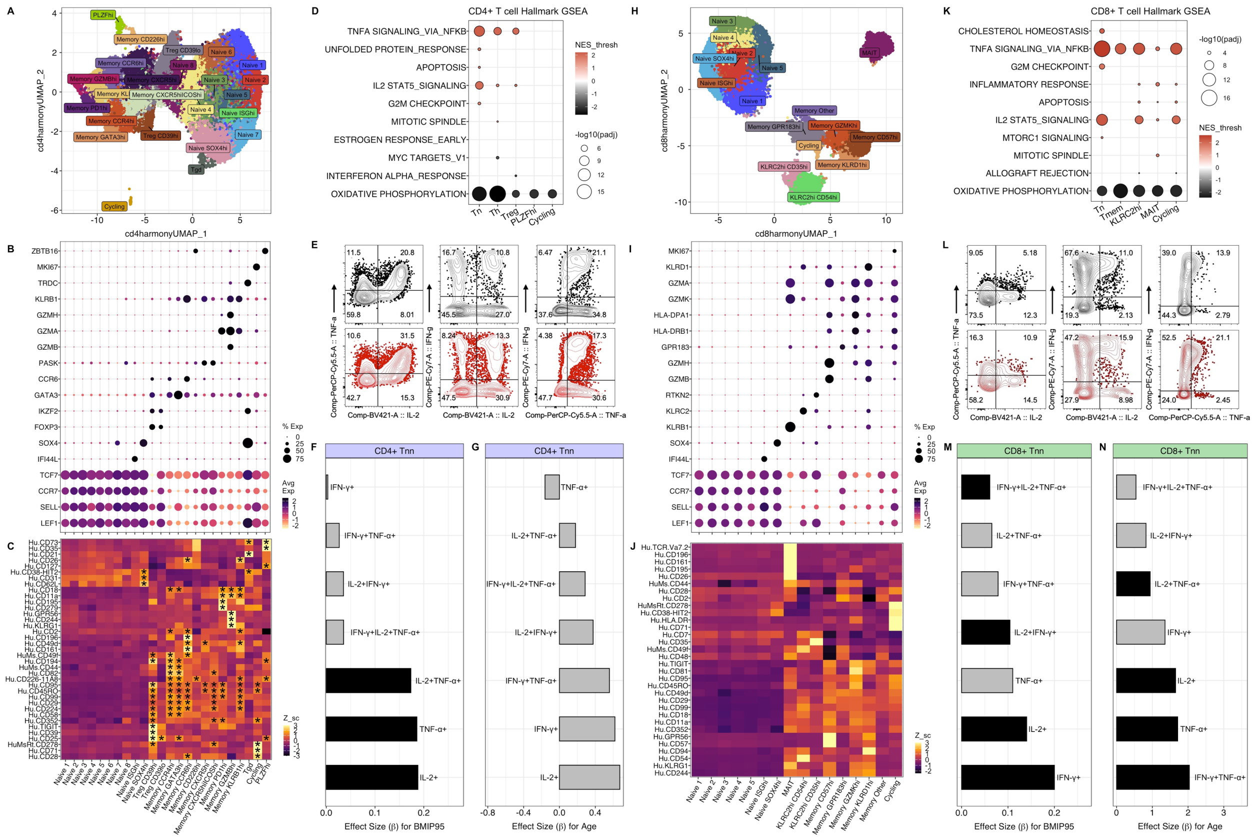

A
B
C
D
E
F
G
H
I
J
K
L
M
N
cd4harmonyUMAP_1
cd4harmonyUMAP_2
CD4+ T cell Hallmark GSEA
TNFA SIGNALING_VIA_NFKB
UNFOLDED PROTEIN_RESPONSE
APOPTOSIS
IL2 STAT5_SIGNALING
G2M CHECKPOINT
MITOTIC SPINDLE
ESTROGEN RESPONSE_EARLY
MYC TARGETS_V1
INTERFERON ALPHA_RESPONSE
OXIDATIVE PHOSPHORYLATION
NES_thresh
-log10(padj)
Tn
Th
Treg
PLZFhi
Cycling
% Exp
Avg Exp
Z_sc
CD4+ Tnn
Effect Size (β) for BMIP95
Effect Size (β) for Age
IFN-γ+
IFN-γ+TNF-α+
IL-2+IFN-γ+
IFN-γ+IL-2+TNF-α+
IL-2+TNF-α+
TNF-α+
IL-2+
Comp-BV421-A :: IL-2
Comp-PerCP-Cy5.5-A :: TNF-a
Comp-PE-Cy7-A :: IFN-g
cd8harmonyUMAP_1
cd8harmonyUMAP_2
CD8+ T cell Hallmark GSEA
CHOLESTEROL HOMEOSTASIS
TNFA SIGNALING_VIA_NFKB
G2M CHECKPOINT
INFLAMMATORY RESPONSE
APOPTOSIS
IL2 STAT5_SIGNALING
MTORC1 SIGNALING
MITOTIC SPINDLE
ALLOGRAFT REJECTION
OXIDATIVE PHOSPHORYLATION
Tn
Tmem
KLRC2hi
MAIT
Cycling
CD8+ Tnn

**Supplemental Figure 2. CITE-seq captures transcriptional dynamics across circulating T cells in pediatric obesity**. Related to Fig. 2. **(A)** UMAP visualization representing 24 distinct $CD4^+$ T cell clusters. **(B)** Bubble plot of transcripts used to distinguish $CD4^+$ T cell clusters. **(C)** Heatmap of selected cell surface protein data as measured by antibody derived tags (ADT) expression. **(D)** Bubble plot of Hallmark gensets across grouped $CD4^+$ T cell clusters differentially enriched (p.adj < 0.001) between HC (black) and O (red) participants. **(E)** Representative gating of indicated cytokines within the $CD4^+$ Tnn compartment following PMA/Ionomycin stimulation. **(F)** Bar plot representing the β-coefficients for the BMIP95 term (Frequency*BMIP95) and **(G)** age term (Frequency*Age) in each model. **(H)** UMAP visualization representing 16 distinct $CD8^+$ T cell clusters. **(I)** Bubble plot of transcripts used to distinguish $CD8^+$ T cell clusters. **(J)** Heatmap of selected ADT expression. **(K)** Bubble plot of Hallmark gensets significantly different (p.adj < 0.001) between HC and O CD8+ T cell cluster groups. **(L)** Representative gating of indicated cytokines within the $CD8^+$ Tnn compartment following PMA/Ionomycin stimulation. **(M)** Bar plot representing the β-coefficients for the BMIP95 term (Frequency*BMIP95) and **(N)** age term (Frequency*Age) in each model. In bubble plots **(B,I),** color scheme reflects average gene expression (scaled log-normalized counts) ranging from -2 (yellow) to 2 (black) and size reflects proportion of cells expressing indicated transcript. In bubble plots **(D,K),** color scheme reflects normalized enrichment score (NES) with negative values in black indicating enrichment in HC and positive values in red indicating enrichment in O and size reflects -Log10 of adjusted p-value. For all heatmaps **(C,J)**, color scheme reflects scaled average area under the curve (AUC) scores across indicated clusters ranging from -3 (black) to 3 (yellow) and cells labeled with an asterisk (*) represent transcripts enriched within the indicated cluster (Wilcoxon's rank-sum test, AUC > 0.75). For all bar plots **(F-G, M-N)** black bars represent significant p-values for indicated model.

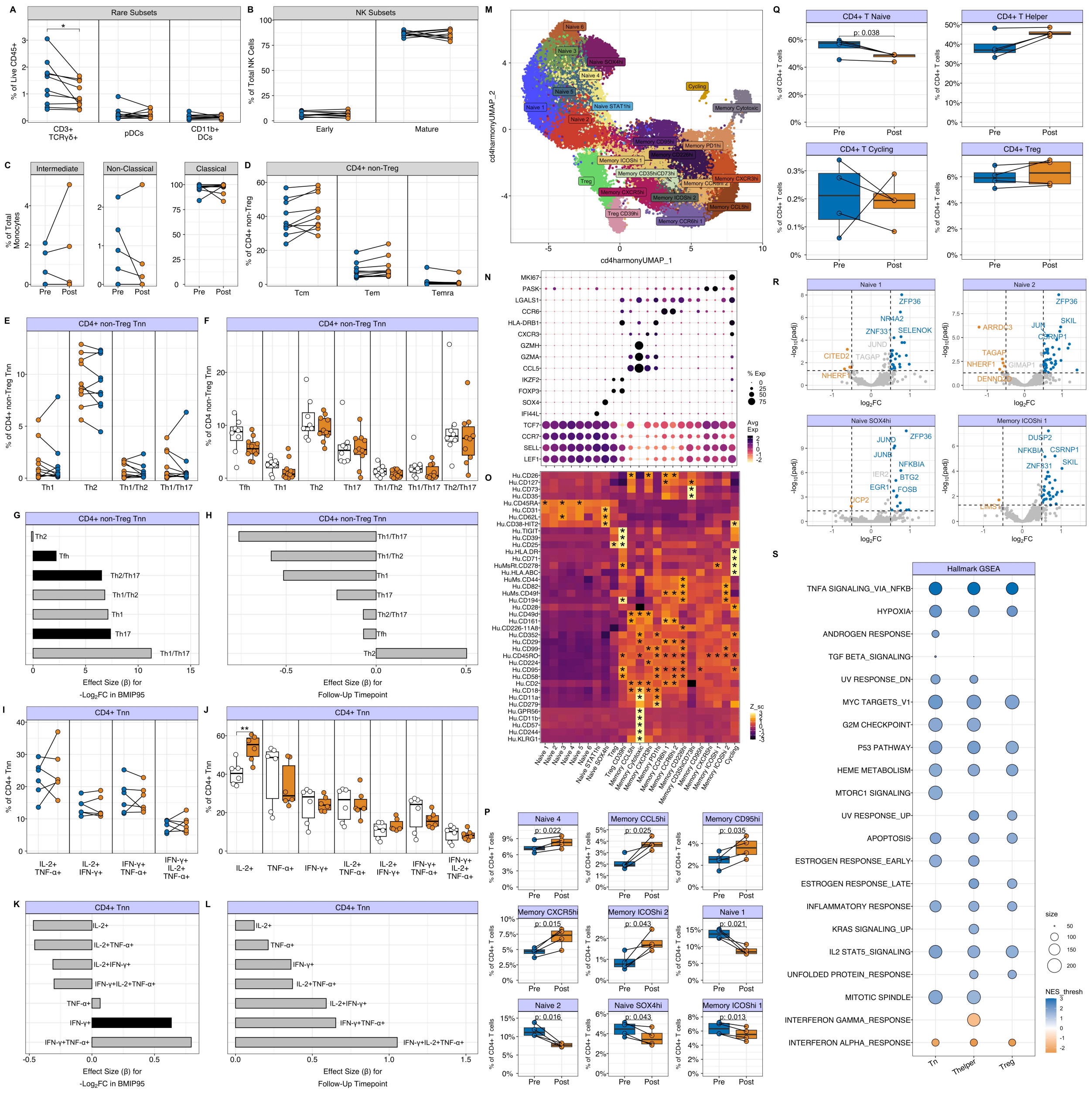
A
Rare Subsets
% of Live CD45+
CD3+ TCRγδ+
pDCs
CD11b+ DCs
B
NK Subsets
% of Total NK Cells
Early
Mature
C
Intermediate
Non-Classical
Classical
% of Total Monocytes
Pre
Post
D
CD4+ non-Treg
% of CD4+ non-Treg
Tcm
Tem
Temra
E
CD4+ non-Treg Tnn
% of CD4+ non-Treg Tnn
Th1
Th2
Th1/Th2
Th1/Th17
F
CD4+ non-Treg Tnn
% of CD4 non-Treg Tnn
Tfh
Th1
Th2
Th17
Th1/Th2
Th1/Th17
Th2/Th17
G
CD4+ non-Treg Tnn
Th2
Tfh
Th2/Th17
Th1/Th2
Th1
Th17
Th1/Th17
Effect Size (β) for -Log2FC in BMIP95
H
CD4+ non-Treg Tnn
Th1/Th17
Th1/Th2
Th1
Th17
Th2/Th17
Tfh
Th2
Effect Size (β) for Follow-Up Timepoint
I
CD4+ Tnn
% of CD4+ Tnn
IL-2+ TNF-α+
IL-2+ IFN-γ+
IFN-γ+ TNF-α+
IFN-γ+ IL-2+ TNF-α+
J
CD4+ Tnn
% of CD4+ Tnn
IL-2+
TNF-α+
IFN-γ+
IL-2+ TNF-α+
IL-2+ IFN-γ+
IFN-γ+ TNF-α+
IFN-γ+ IL-2+ TNF-α+
K
CD4+ Tnn
IL-2+
IL-2+TNF-α+
IL-2+IFN-γ+
IFN-γ+IL-2+TNF-α+
TNF-α+
IFN-γ+
IFN-γ+TNF-α+
Effect Size (β) for -Log2FC in BMIP95
L
CD4+ Tnn
IL-2+
TNF-α+
IFN-γ+
IL-2+TNF-α+
IL-2+IFN-γ+
IFN-γ+TNF-α+
IFN-γ+IL-2+TNF-α+
Effect Size (β) for Follow-Up Timepoint
M
cd4harmonyUMAP_2
cd4harmonyUMAP_1
Naive 6
Naive 3
Naive SOX4hi
Naive 4
Naive 5
Naive 1
Naive STAT1hi
Naive 2
Cycling
Memory Cytotoxic
Memory CD95hi
Memory PD1hi
Memory ICOShi 1
Memory CD35hiCD73hi
Memory CXCR3hi
Treg
Memory CXCR5hi
Memory ICOShi 2
Memory CCL5hi
Treg CD39hi
Memory CCR6hi 1
N
MKI67
PASK
LGALS1
CCR6
HLA-DRB1
CXCR3
GZMH
GZMA
CCL5
IKZF2
FOXP3
SOX4
IFI44L
TCF7
CCR7
SELL
LEF1
% Exp
Avg Exp
O
Z_sc
Q
CD4+ T Naive
p: 0.038
CD4+ T Helper
CD4+ T Cycling
CD4+ Treg
% of CD4+ T cells
Pre
Post
R
Naive 1
ZFP36
NR4A2
ZNF331
SELENOK
JUND
CITED2
TAGAP
NHERF1
Naive 2
ARRDC3
JUN
CSRNP1
SKIL
TAGAP
NHERF1
GIMAP1
DENND4A
Naive SOX4hi
JUND
ZFP36
JUNB
NFKBIA
IER2
BTG2
EGR1
FOSB
DCP2
Memory ICOShi 1
DUSP2
NFKBIA
CSRNP1
ZNF331
SKIL
-log10(padj)
log2FC
P
Naive 4
p: 0.022
Memory CCL5hi
p: 0.025
Memory CD95hi
p: 0.035
Memory CXCR5hi
p: 0.015
Memory ICOShi 2
p: 0.043
Naive 1
p: 0.021
Naive 2
p: 0.016
Naive SOX4hi
p: 0.043
Memory ICOShi 1
p: 0.013
% of CD4+ T cells
Pre
Post
S
Hallmark GSEA
TNFA SIGNALING_VIA_NFKB
HYPOXIA
ANDROGEN RESPONSE
TGF BETA_SIGNALING
UV RESPONSE_DN
MYC TARGETS_V1
G2M CHECKPOINT
P53 PATHWAY
HEME METABOLISM
MTORC1 SIGNALING
UV RESPONSE_UP
APOPTOSIS
ESTROGEN RESPONSE_EARLY
ESTROGEN RESPONSE_LATE
INFLAMMATORY RESPONSE
KRAS SIGNALING_UP
IL2 STAT5_SIGNALING
UNFOLDED PROTEIN_RESPONSE
MITOTIC SPINDLE
INTERFERON GAMMA_RESPONSE
INTERFERON ALPHA_RESPONSE
Tn
Thelper
Treg
size
NES_thresh

**Supplemental Figure 3. Multimodal analysis reveals functional and transcriptional alterations in peripheral CD4$^+$ T cells following weight loss.** Related to Fig.3.

**(A)** Indicated population frequencies expressed as proportion of live CD45$^+$ cells. **(B)** NK cell subsets shown as frequency of total NK cells. **(C)** Monocyte subsets shown as frequency of total monocytes. **(D)** Frequency of indicated memory subsets within CD4$^+$ Teff compartment. **(E-F)** Frequency of indicated T helper subset shown as proportion of non-naive CD4$^+$ Teff cells across **(E)** paired bariatric surgery participants **(F)** and between HC (white) and Post participants. **(G)** Bar plot representing the β-coefficients for the -Log2FC BMIP95 term (Log2FCFrequency *- Log2 BMIP95) and **(H)** follow-up visit term (Log2FCFrequency * months since surgery) in each model. **(I-J)** Frequency of indicated cytokine within CD4$^+$ Tnn compartment across **(I)** paired bariatric surgery participants and **(J)** across HC (white) and Post participants. **(K)** Bar plot representing the β-coefficients for the -Log2FC BMIP95 term (Log2FCFrequency *- Log2 BMIP95) and **(L)** follow-up visit term (Log2FCFrequency * months since surgery) in each model. **(M)** UMAP visualization representing 24 distinct CD4$^+$ T cell clusters. **(N)** Bubble plot of transcripts used to distinguish CD4$^+$ T cell clusters. Color scheme reflects average gene expression (scaled log-normalized counts) ranging from -2 (yellow) to 2 (black) and size reflects proportion of cells expressing indicated transcript. **(O)** Heatmap of selected cell surface protein data as measured by ADT expression. Color scheme reflects scaled average AUC scores across indicated clusters ranging from -3 (black) to 3 (yellow). Cells labeled with an asterisk (*) represent transcripts enriched within the indicated cluster (Wilcoxon's rank-sum test, AUC > 0.75). **(P-Q)** Cluster abundance in paired Pre and Post bariatric surgery samples across indicated **(P)** unique clusters and **(Q)** cluster groups. **(R)** Volcano plot comparing DEGs across indicated clusters. Vertical dashed lines indicate absolute value of log2FC in expression >0.5 and horizontal dashed line represents significance cut off (p< 0.05). Dots represent individual transcripts with blue indicating transcripts enriched in Pre and orange representing transcripts enriched in Post. Gray represents DEGs that did not meet log2FC or significance cutoffs. **(S)** Bubble plot of Hallmark gensets different (p.adj < 0.001) between Pre and Post CD4$^+$ T cell cluster groups. Color scheme reflects NES values with blue (positive NES) indicating enrichment in Pre and orange (negative NES) indicating enrichment in Post and size reflects -Log10 of adjusted p-value. In bar plots **(G-H, K-L)** black bars represent significant p-values (<0.05) for indicated model. For all boxplots and line points, circles represent individual donors across HC (white), Pre (blue) and Post (orange) participants with matched donors connected by a line. Paired donors evaluated via paired t-test whereas HC vs Post comparisons evaluated by pairwise t-test ($* = p < 0.05$, $** = p < 0.01$, $*** = p < 0.001$).

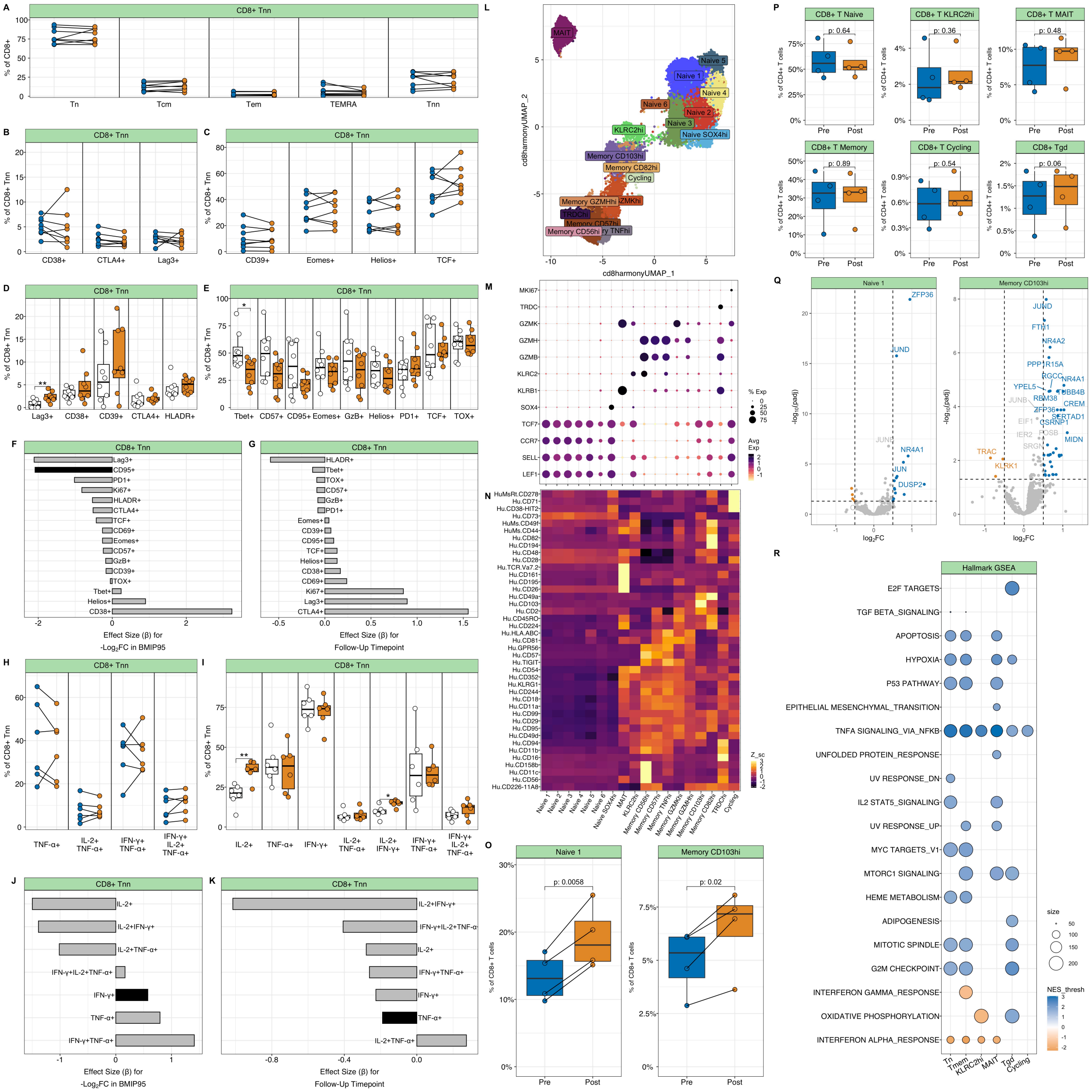

A
CD8+ Tnn
% of CD8+
Tn
Tcm
Tem
TEMRA
Tnn
B
C
% of CD8+ Tnn
CD38+
CTLA4+
Lag3+
CD39+
Eomes+
Helios+
TCF+
D
E
HLADR+
Tbet+
CD57+
CD95+
GzB+
PD1+
TOX+
F
G
Effect Size (β) for
-Log2FC in BMIP95
Follow-Up Timepoint
Ki67+
CD69+
H
I
TNF-α+
IL-2+ TNF-α+
IFN-γ+ TNF-α+
IFN-γ+ IL-2+ TNF-α+
IL-2+
IFN-γ+
IL-2+ IFN-γ+
J
K
IL-2+IFN-γ+
IL-2+TNF-α+
IFN-γ+IL-2+TNF-α+
IFN-γ+TNF-α+
L
cd8harmonyUMAP_1
cd8harmonyUMAP_2
MAIT
Naive 1
Naive 2
Naive 3
Naive 4
Naive 5
Naive 6
Naive SOX4hi
KLRC2hi
Memory CD103hi
Memory CD82hi
Cycling
Memory GZMHhi
GZMKhi
TRDChi
Memory CD57hi
Memory CD56hi
ry TNFhi
M
MKI67
TRDC
GZMK
GZMH
GZMB
KLRC2
KLRB1
SOX4
TCF7
CCR7
SELL
LEF1
% Exp
Avg Exp
N
Z_sc
O
Naive 1
Memory CD103hi
% of CD8+ T cells
p: 0.0058
p: 0.02
Pre
Post
P
CD8+ T Naive
CD8+ T KLRC2hi
CD8+ T MAIT
CD8+ T Memory
CD8+ T Cycling
CD8+ Tgd
% of CD4+ T cells
p: 0.64
p: 0.36
p: 0.48
p: 0.89
p: 0.54
p: 0.06
Q
Naive 1
Memory CD103hi
-log10(padj)
log2FC
ZFP36
JUND
NR4A1
JUN
DUSP2
FTH1
NR4A2
CREM
MIDN
TRAC
KLRK1
R
Hallmark GSEA
E2F TARGETS
TGF BETA_SIGNALING
APOPTOSIS
HYPOXIA
P53 PATHWAY
EPITHELIAL MESENCHYMAL_TRANSITION
TNFA SIGNALING_VIA_NFKB
UNFOLDED PROTEIN_RESPONSE
UV RESPONSE_DN
IL2 STAT5_SIGNALING
UV RESPONSE_UP
MYC TARGETS_V1
MTORC1 SIGNALING
HEME METABOLISM
ADIPOGENESIS
MITOTIC SPINDLE
G2M CHECKPOINT
INTERFERON GAMMA_RESPONSE
OXIDATIVE PHOSPHORYLATION
INTERFERON ALPHA_RESPONSE
size
NES_thresh
Tn
Tmem
KLRC2hi
MAIT
Tgd
Cycling

**Supplemental Figure 4. Altered transcriptional landscape of peripheral CD8$^+$ T cells following weight loss.** Related to Fig.4. **(A)** Frequency of indicated memory subsets within CD8$^+$ T cells. **(B-D)** Frequency of indicated populations within CD8$^+$ Tnn compartment across paired bariatric surgery **(B-C)** and between Pre and HC **(D-E)**. **(F)** Bar plot representing the β-coefficients for the -Log2FC BMIP95 term (Log2FCFrequency *- Log2 BMIP95) and **(G)** follow-up visit term (Log2FCFrequency * months since surgery) in each model. **(H-I)** Frequency of indicated cytokine within CD8$^+$ Tnn cells between paired bariatric surgery participants **(H)** and between Pre and HC **(I). (J)** Bar plot representing the β-coefficients for the -Log2FC BMIP95 term (Log2FCFrequency *- Log2 BMIP95) and **(K)** follow-up visit term (Log2FCFrequency * months since surgery) in each model. **(K)** Frequency of indicated cytokine within CD8$^+$ Tnn cells. **(L)** UMAP visualization representing 28 distinct CD8$^+$ T cell clusters. **(M)** Bubble plot of transcripts used to distinguish CD8$^+$ T cell clusters. Color scheme reflects average gene expression (scaled log-normalized counts) ranging from -2 (yellow) to 2 (black) and size reflects proportion of cells expressing indicated transcript. **(N)** Heatmap of selected cell surface protein data as measured by ADT expression. Color scheme reflects scaled average AUC scores across indicated clusters ranging from -3 (black) to 3 (yellow). **(O-P)** Cluster abundance in paired Pre and Post bariatric surgery samples across indicated **(O)** unique clusters and **(P)** cluster groups. **(Q)** Volcano plot comparing DEGs across indicated clusters. Vertical dashed lines indicate absolute value of log2FC in expression >0.5 and horizontal dashed line represents significance cut off ($p < 0.05$). Dots represent individual transcripts with blue indicating transcripts enriched in Pre and orange representing transcripts enriched in Post. Gray represents DEGs that did not meet log2FC or significance cutoffs. **(R)** Bubble plot of Hallmark gensets different (p.adj < 0.001) between Pre and Post CD8$^+$ T cell cluster groups. Color scheme reflects NES values with blue (positive NES) indicating enrichment in Pre and orange (negative NES) indicating enrichment in Post and size reflects -Log10 of adjusted p-value. In bar plots **(G-H, J-K)** black bars represent significant p-values (<0.05) for indicated model. For all boxplots and line points, circles represent individual donors across HC (white), Pre (blue) and Post (orange) participants with matched donors connected by a line. Paired donors evaluated via paired t-test whereas HC vs Post comparisons evaluated by pairwise t-test ($* = p < 0.05$, $** = p < 0.01$, $*** = p < 0.001$).
